# A potently neutralizing anti-SARS-CoV-2 antibody inhibits variants of concern by binding a highly conserved epitope

**DOI:** 10.1101/2021.04.26.441501

**Authors:** Laura A. VanBlargan, Lucas J. Adams, Zhuoming Liu, Rita E. Chen, Pavlo Gilchuk, Saravanan Raju, Brittany K. Smith, Haiyan Zhao, James Brett Case, Emma S. Winkler, Bradley M. Whitener, Lindsay Droit, Ishmael D. Aziati, Pei-Yong Shi, Adrian Creanga, Amarendra Pegu, Scott A. Handley, David Wang, Adrianus C.M. Boon, James E. Crowe, Sean P.J. Whelan, Daved H. Fremont, Michael S. Diamond

## Abstract

With the emergence of SARS-CoV-2 variants with increased transmissibility and potential resistance, antibodies and vaccines with broadly inhibitory activity are needed. Here we developed a panel of neutralizing anti-SARS-CoV-2 mAbs that bind the receptor binding domain of the spike protein at distinct epitopes and block virus attachment to cells and its receptor, human angiotensin converting enzyme-2 (hACE2). While several potently neutralizing mAbs protected K18-hACE2 transgenic mice against infection caused by historical SARS-CoV-2 strains, others induced escape variants *in vivo* and lost activity against emerging strains. We identified one mAb, SARS2-38, that potently neutralizes all SARS-CoV-2 variants of concern tested and protects mice against challenge by multiple SARS-CoV-2 strains. Structural analysis showed that SARS2-38 engages a conserved epitope proximal to the receptor binding motif. Thus, treatment with or induction of inhibitory antibodies that bind conserved spike epitopes may limit the loss of potency of therapies or vaccines against emerging SARS-CoV-2 variants.

## INTRODUCTION

Severe acute respiratory syndrome-related coronavirus (SARS-CoV) and SARS-CoV-2 belong to the Sarbecovirus subgenus of Betacoronaviruses (Viruses, 2020). In little more than a year, the coronavirus disease 2019 (COVID-19) pandemic caused by the rapid emergence of SARS-CoV-2 has resulted in over 140 million infections and 3 million deaths worldwide (https://covid19.who.int/). Multiple effective vaccines against SARS-CoV-2 that prevent COVID-19 have been rapidly developed and deployed (Baden et al., 2021; Polack et al., 2020; Sadoff et al., 2021; Voysey et al., 2021). Monoclonal antibodies (mAb) also have shown efficacy in animal models of SARS-CoV-2 infection (Alsoussi et al., 2020; Baum et al., 2020a; Fagre et al., 2020; Hansen et al., 2020; Hassan et al., 2020; Kreye et al., 2020; Rogers et al., 2020; Shi et al., 2020; Zost et al., 2020), and two mAb treatments are approved for use in patients under Emergency Use Authorization (EUA) (Chen et al., 2021b; Weinreich et al., 2021). Therapy with mAbs may be beneficial to high-risk patients following exposure to SARS-CoV-2 with mild or moderate symptoms, but prior to onset of severe disease signs and symptoms, and can complement the usage of vaccines as a means of combating the COVID-19 pandemic.

The majority of characterized potently neutralizing and protective anti-SARS-CoV-2 mAbs bind the receptor binding domain (RBD) of the viral spike protein (Barnes et al., 2020; Baum *et al*., 2020a; Cao et al., 2020; Tortorici et al., 2020; Zost *et al*., 2020), though some inhibitory mAbs against the N-terminal domain (NTD) of spike also have been described (Chi et al., 2020; Liu et al., 2020; Suryadevara et al., 2021). Under immune selection pressure, SARS-CoV-2 can select for mutations in the RBD and NTD that enable escape from antibody recognition and neutralization (Baum et al., 2020b; Greaney et al., 2021; Liu et al., 2021; Starr et al., 2021; Suryadevara *et al*., 2021). Indeed, several emerging SARS-CoV-2 variants have mutations in the spike protein, including the RBD and NTD, that confer resistance to mAbs or polyclonal antibodies (pAbs) elicited by vaccines or natural infection (Chen et al., 2021d; Thomson et al., 2021; Weisblum et al., 2020). As such, additional mAbs or vaccines that retain efficacy against emerging SARS-CoV-2 variants may be needed to combat new and evolving strains.

In this study, we describe a panel of potently neutralizing murine mAbs against the RBD of SARS-CoV-2 that bind several epitopes proximal to the receptor binding motif (RBM) of the RBD or at the base of the RBD. Although some neutralizing mAbs demonstrated limited ability to protect against infection by the historical SARS-CoV-2 WA1/2020 strain in a mouse disease model and selected for rapid escape *in vivo*, others protected completely in the context of prophylactic or therapeutic administration. Two protective mAbs, SARS2-02 and SARS2-38, showed variable capacity to neutralize variants of concern (VOCs): SARS2-02 binds an epitope that includes residues E484 and L452 and has reduced potency against strains (B.1.429, B.1.351, and B.1.1.28) encoding these mutations. In contrast, SARS2-38 binds an epitope centered on residues K444 and G446 and potently neutralized all tested VOCs. Analysis of a cryo-electron microscopy (cryo-EM) structure of SARS2-38 bound to spike reveals that this mAb binds a conserved epitope on the RBD that is also engaged, albeit through distinct geometries, by other neutralizing and protective human mAbs. Thus, treatment with mAbs or induction of pAbs targeting this conserved region of the RBD may confer protection against many emerging SARS-CoV-2 variants.

## RESULTS

### Development and characterization of anti-SARS-CoV-2 mAbs

We generated a panel of anti-SARS-CoV-2 mAbs from BALB/c mice that were immunized and boosted with purified RBD and/or ectodomain of the spike protein mixed with AddaVax™, a squalene-based adjuvant (**Fig 1**). After splenocyte-myeloma fusions, hybridoma supernatants were screened for antibody binding to recombinant spike protein and permeabilized SARS-CoV-2-infected Vero cells by ELISA and flow cytometry, respectively. Sixty-four hybridomas producing anti-SARS-CoV-2 antibodies were cloned by limiting dilution. Forty-three of these mAbs bound to recombinant RBD and were selected for further study because prior experiments showed this class included potently inhibitory antibodies (Barnes *et al*., 2020; Baum *et al*., 2020a; Cao *et al*., 2020; Tortorici *et al*., 2020; Zost *et al*., 2020); the majority of these mAbs were of the IgG1 subclass (**Fig 1**).

**Figure 1.**
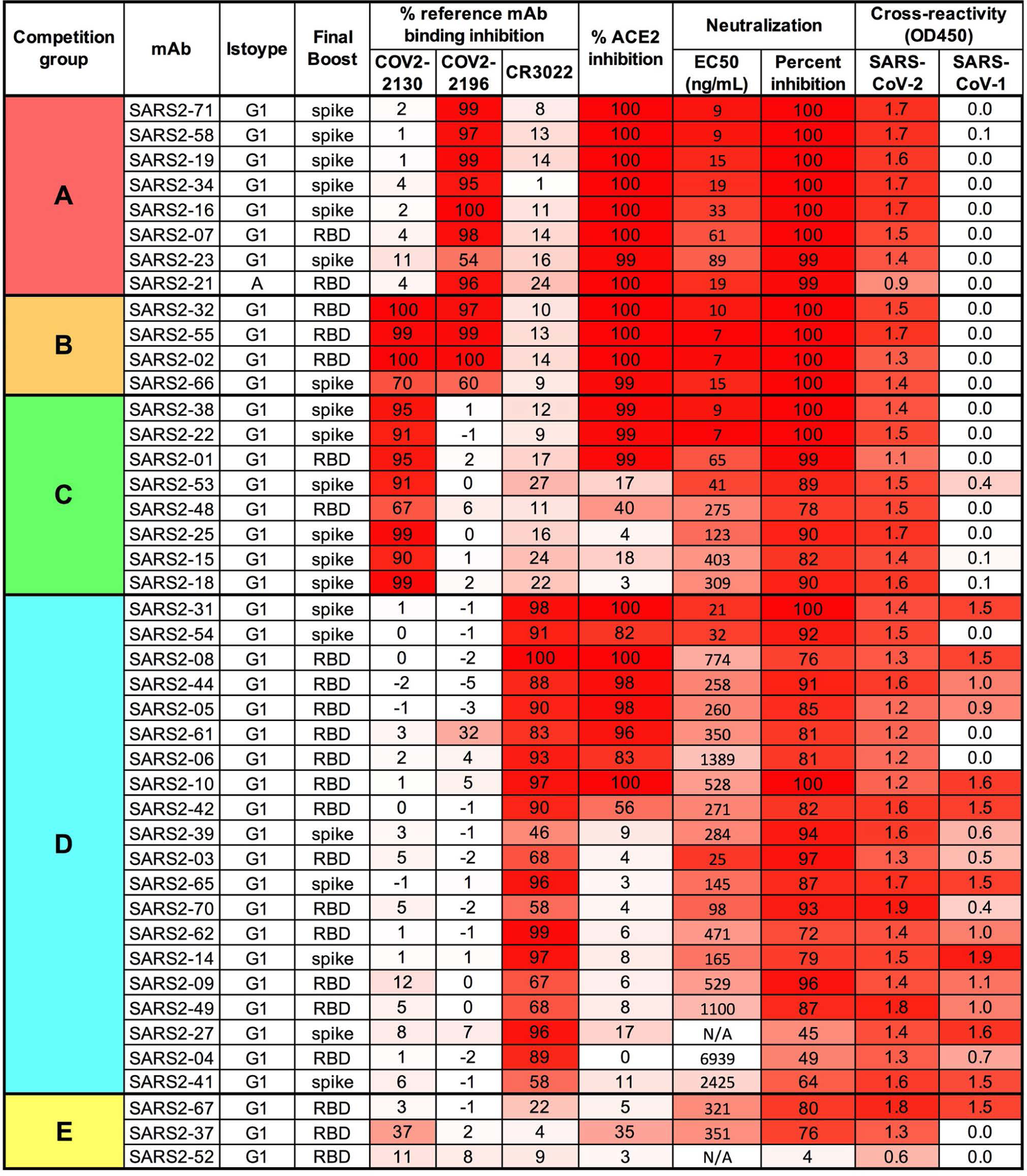
Panel of anti-SARS-CoV-2 mAbs. Hybridoma supernatants from the panel of anti-SARS-CoV-2 murine mAbs were assayed for neutralization of SARS-CoV-2 by FRNT, cross-reactivity to SARS-CoV-1 spike protein, and ability to inhibit SARS-CoV-2 spike protein binding to hACE2 or a panel of reference human mAbs through competition ELISA. MAbs are grouped by reference mAb competition properties. Data represent the mean (or geometric mean for EC50 values) from two to four independent experiments. Hybridomas were produced from splenocytes of mice that received three immunizations (once with the RBD and then twice with Spike) prior to a final pre-fusion boost with either RBD or Spike, as indicated in the ‘Final Boost’ column.

The mAbs were evaluated by competition binding analysis using three previously characterized human mAbs that recognize distinct antigenic sites on the RBD (COV2-2196, COV2-2130, and CR3022) (Yuan et al., 2020; Zost *et al*., 2020) (**Fig 1**). Eight mAbs competed for spike protein binding with the neutralizing mAb COV2-2196 only, eight mAbs competed with the neutralizing mAb COV2-2130 only, four mAbs competed with both COV2-2196 and COV2-2130, and twenty mAbs competed with CR3022, which recognizes a more conserved, cryptic, and non-neutralizing epitope on the SARS-CoV-2 spike protein distal from the receptor binding site. Three RBD-binding mAbs did not compete with COV2-2196, COV2-2130, or CR3022. Based on the binding analysis, mAbs were divided into 5 competition groups, A-E (**Fig 1**).

One potential mechanism of antibody-mediated neutralization of SARS-CoV-2 is through inhibition of viral spike protein binding to the human ACE2 receptor. The COV2-2196 epitope directly overlaps the ACE2 binding site on RBD, whereas the COV2-2130 epitope lies proximal to residues in the RBM that interact with ACE2 (Dong et al., 2021); nonetheless, both mAbs can block spike binding to ACE2. In contrast, CR3022 engages the base of the RBD and does not block ACE2 binding to spike (Yuan *et al*., 2020). Of the 43 RBD-binding antibodies in our panel, all mAbs in groups A and B inhibited ACE2 binding to spike protein, mAbs in groups C and D variably inhibited ACE2 binding, and mAbs in group E failed to inhibit ACE2 binding (**Fig 1**).

The mAbs also were tested for cross-reactive binding to the SARS-CoV-1 spike protein. The majority of mAbs in group D, which competed with the cross-reactive mAb CR3022 for spike binding, cross-reacted with SARS-CoV-1 spike protein, indicating they bind conserved sarbecovirus epitopes. MAbs in groups A, B, and C did not bind to SARS-CoV-1 (**Fig 1**), and one group E mAb recognized SARS-CoV-1.

### Neutralizing activity of anti-SARS-CoV-2 mAbs

We next determined the neutralizing activity of mAb hybridoma supernatants using a focus-reduction neutralization test (FRNT) and Vero E6 cells (Case et al., 2020) with the WA1/2020 SARS-CoV-2 strain. Antibody concentrations in the supernatants were quantified by ELISA and used to calculate half-maximal inhibitory concentrations (EC50 values). The most potently inhibitory mAbs (EC50: < 10 ng/mL) belonged to groups A, B, and C, and also blocked ACE2 binding (**Fig 1**). Some mAbs in group C and D that did not block ACE2 binding still showed robust neutralizing activity (EC50: 20 – 100 ng/mL), although the majority were weakly inhibitory. Group E mAbs were weakly neutralizing and did not block ACE2 binding.

A subset of mAbs from groups A, B, C, and D were selected for more detailed study. We chose two mAbs with the highest neutralization potency from each group; in cases where mAbs had high variable region sequence similarity, we selected only one of these mAbs for further study. We also selected SARS2-03, as it was one of the few neutralizing mAbs that did not block ACE2 binding. Nine mAbs were purified from hybridoma supernatants and retested for neutralization potency by FRNT using Vero cells and the WA1/2020 isolate (**Fig 2A-B**). Again, the most potently neutralizing purified mAbs belonged to groups A, B, and C, with less inhibitory activity in those derived from group D. We also characterized these nine mAbs for competition binding with each other (**Fig S1**). The two group A mAbs (SARS2-34 and SARS2-71) competed for spike binding only with each other. In contrast, mAbs in groups B (SARS2-02 and SARS2-55) and C (SARS2-01 and SARS2-38) competed for spike binding across both groups. SARS2-03, a group D mAb, did not bind spike efficiently in the presence of group B or C mAbs and blocked binding of group C mAb SARS2-01. SARS2-10 and SARS2-31, the other two group D mAbs, however, competed with only each other. Together, these results suggest that mAbs in group C may have overlapping epitopes with group B mAbs and group D mAb SARS2-03, whereas group A mAbs and the remaining group D mAbs likely engage physically distinct epitopes.

**Figure 2.**
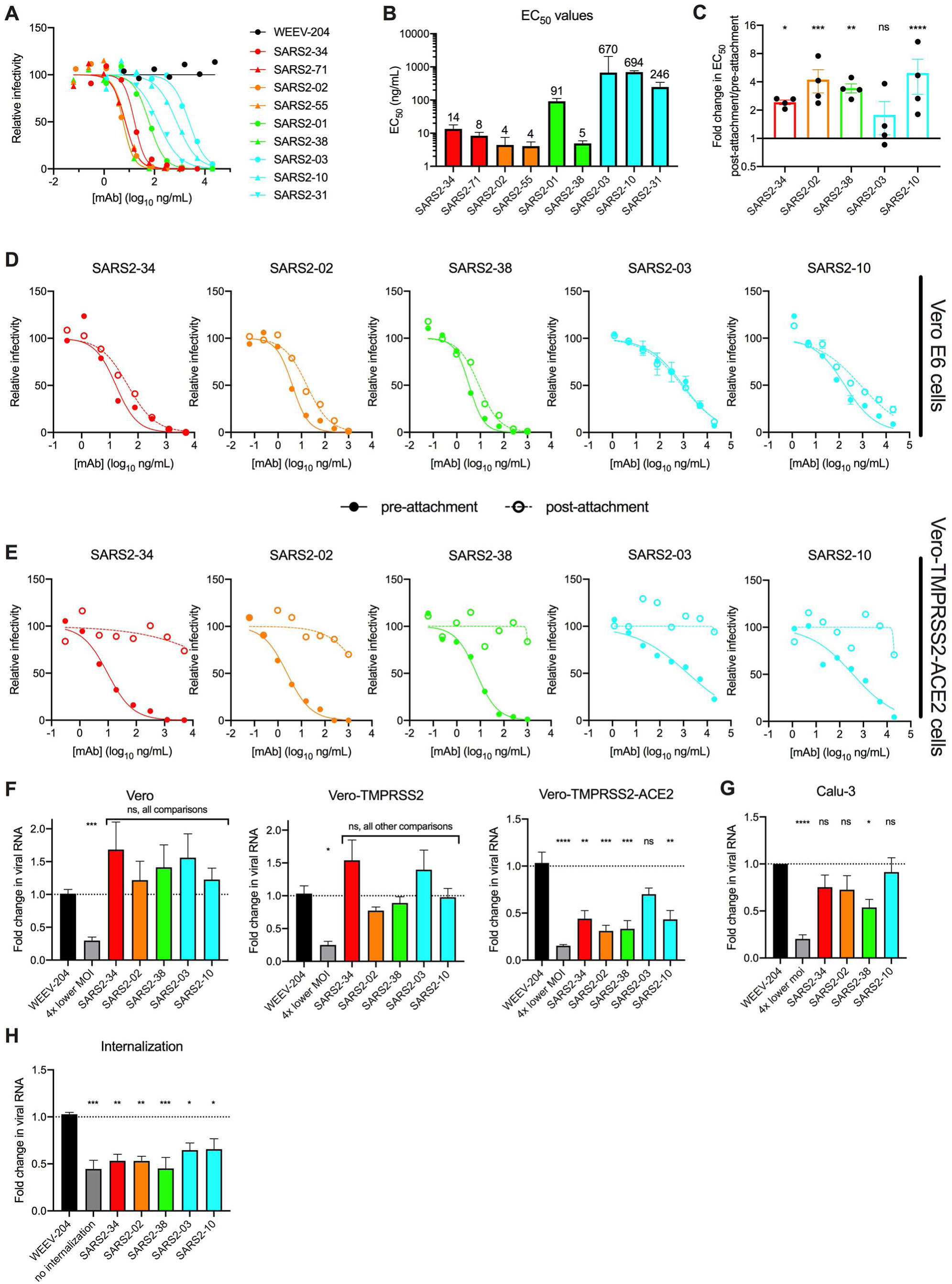
Neutralization by anti-SARS-CoV-2 mAbs. (**A-B**) Anti-SARS-CoV-2 mAbs were assayed for neutralization by FRNT against SARS-CoV-2 using Vero E6 cells. (**A**) Representative dose response curves are shown. (**B**) Mean EC50 values are shown; data are from three to four experiments. (**C-D**) Anti-SARS-CoV-2 mAbs were assayed for pre- or post-attachment neutralization of SARS-CoV-2 using Vero E6 cells. (**C**) Fold change in EC50 values for post-attachment over pre-attachment neutralization. Error bars represent standard error of the mean (SEM) from four experiments (**D**) Representative dose response curves are shown. (**E**) Anti-SARS-CoV-2 mAbs were assayed for pre- or post-attachment inhibition on Vero-TMPRSS2-ACE2 cells. Dose response curves are shown. Data are representative of three experiments. (**F-G**) Anti-SARS-CoV-2 mAbs were assayed for attachment inhibition of SARS-CoV-2 to Vero E6, Vero-TMPRSS2, or Vero-TMPRSS2-ACE2 (**F**) or Calu-3 (**G**) cells. Data are from three (**F**) to six (**G**) experiments. (**H**) Anti-SARS-CoV-2 mAbs were assayed for inhibition of virus internalization in Vero E6 cells. Data are from four experiments. **C**. ANOVA with Sidak’s post-test comparing pre- vs. post-attachment EC50 values for each mAb; **F-H**. One-way ANOVA with Dunnett’s post-test compared mAb treatment to isotype control mAb treatment. ns, not significant; *p<0.05; **p<0.01; ***p<0.001, ****p<0.0001.

### Mechanism of neutralization by anti-SARS-CoV-2 mAbs

We investigated whether the anti-SARS-CoV-2 mAbs inhibited infection at a pre- or post-attachment step of the entry process. For these experiments, we selected one representative mAb from groups A, B, and C (SARS2-34, SARS2-02, and SARS2-38, respectively) and two mAbs from group D (one that blocks ACE2 binding, [SARS2-10] and one that does not [SARS2-03]). We compared the neutralization potency of mAbs when added before or after virus absorption to Vero E6 cells. Unexpectedly, all mAbs retained neutralizing activity when added post-attachment, although the potency of groups A, B, and C mAbs SARS2-02, SARS2-34, and SARS2-38 was reduced slightly (∼2- to 4-fold, p < 0.05) relative to pre-attachment neutralization titers (**Fig 2C-D**). SARS2-10, a group E mAb, also showed a ∼5-fold decrease (p < 0.0001) in neutralizing activity when added after attachment. In contrast, SARS2-03, another group E mAb, and the only mAb in this smaller panel that did not block ACE2-spike interactions, had similar neutralization potencies (p = 0.79) when added before or after cell attachment. These data suggest that mAbs that inhibit spike protein binding to ACE2 neutralize SARS-CoV-2 slightly more efficiently when given at a pre-attachment step, although all of the mAbs tested retained the ability to inhibit infection when given after virus attachment to cells.

To determine the impact of entry factor expression on target cells on virus neutralization, we extended these findings to cells that ectopically express human ACE2 and TMPRSS2. In contrast to the relatively minor change in neutralization potency seen with all mAbs for pre- versus post-attachment observed using Vero E6 cells, mAbs no longer efficiently neutralized SARS-CoV-2 infection when added after attachment to Vero-TMPRSS2-ACE2 cells, although pre-attachment neutralization activity remained intact (**Fig 2E**). Thus, the ability of anti-SARS-CoV-2 mAbs to neutralize at a post-attachment step depended on expression levels of viral entry factors and the cell line.

We also tested the ability of the mAbs to block directly virus attachment to cells, including Vero E6, Vero-TMPRSS2, and Vero-TMPRSS2-ACE2 cells. None of the mAbs efficiently blocked SARS-CoV-2 attachment to Vero or Vero-TMPRSS2 cells (**Fig 2F**). However, with the exception of SARS2-03, all mAbs reduced virus attachment to Vero-TMPRSS2-ACE2. To corroborate these findings with cells that endogenously express human ACE2, we repeated experiments with Calu-3 cells, a human lung epithelial cell line. We observed an intermediate phenotype with Calu-3 cells, with modest attachment inhibition by mAbs in groups A, B, and C; levels of attached virus were ∼25-50% lower than the isotype mAb control with inhibition by only SARS2-38 attaining statistical significance (**Fig 2G**). This result suggests that the anti-RBD mAbs can inhibit viral attachment to cells, but this activity depends on levels of human ACE2 expression. Since the mAbs did not efficiently inhibit attachment to Vero E6 cells lacking human ACE2 expression, we tested whether they block a later step in the entry process by using a virus internalization assay (Dejarnac et al., 2018; Earnest et al., 2021). In Vero E6 cells, pre-incubation with all of the anti-RBD mAbs tested resulted in reduced levels of internalized virus (**Fig 2H**).

Because we observed cell type-dependent differences in the mechanism of neutralization, we tested the effect of cell substrate on the inhibitory potency of our anti-RBD mAbs by FRNT. Notably, the anti-RBD mAbs neutralized SARS-CoV-2 WA1/2020 equivalently in Vero E6, Vero-TMRPSS2, and Vero-TMPRSS2-ACE2 cells (**Fig S2**). Thus, although the mAbs variably block SARS-CoV-2 attachment on different cell types, the potency of infection inhibition was similar across cell substrates. This result may be explained by the ability of anti-RBD mAbs to block a required ACE2-dependent entry interaction in all of the cell substrates tested, even though the attachment step is variably affected.

### Epitope mapping of anti-SARS-CoV-2 mAbs using neutralization escape analysis

To determine spike residues important for recognition by anti-SARS-CoV-2 mAbs, we previously isolated neutralization escape mutants by passaging a VSV-eGFP-SARS-CoV-2-S chimeric virus in the presence of neutralizing mAbs, including some of the antibodies described in this study (Liu *et al*., 2021). The above described subset of nine mAbs from groups A-D were tested for neutralization against the panel of sequenced mutants, with the exception of SARS2-10 and SARS2-03, which were not evaluated because of difficultly in isolating escape mutants with mAbs of low neutralization potency. Neutralizing activity was lost for group A mAbs SARS2-34 and SARS2-71 when residues 476-479, 486, and 499 were mutated; for group B mAbs SARS2-02 and SARS2-55 when residues 446, 452, and 484 were mutated; for group C mAb SARS2-01 when residues 346, 352, 446, 450, and 494 were mutated; for group C mAb SARS2-38 when residues 444 and 446 were mutated; and for group D mAb SARS2-31 when residues 378, 408, and 504 were mutated (**Fig 3A-F**).

**Figure 3.**
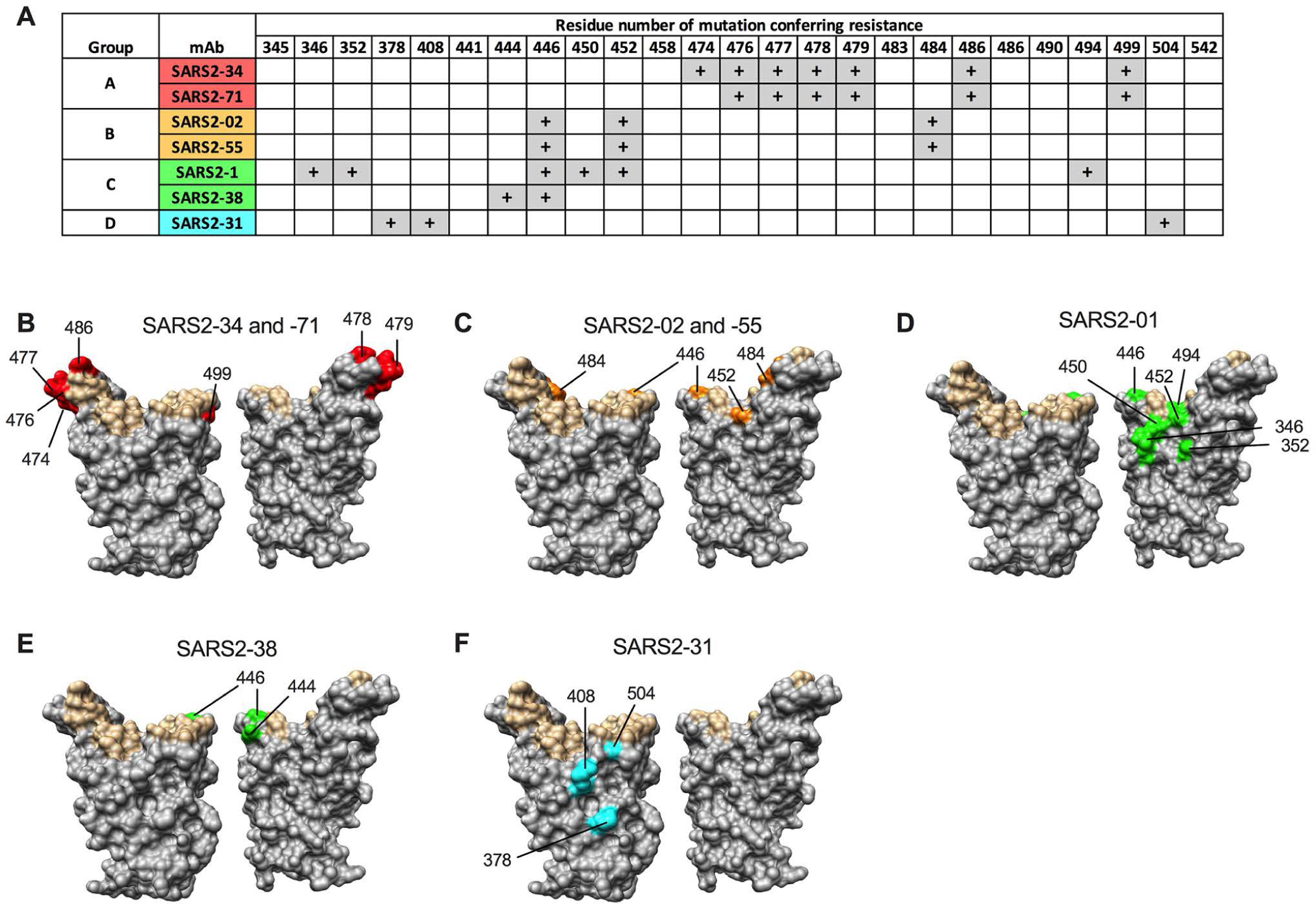
Epitopes recognized by anti-SARS-CoV-2 mAbs. mAbs were tested for neutralization potency against a panel of VSV-eGFP-SARS-COV-2-S neutralization escape mutants. (**A**) “+” symbol indicates resistance to neutralization when a mutation at the indicated residue number is present. (**B-F**) Residues from (**A**) are highlighted on the RBD structure (PDB 6M0J) in red, orange, green, or cyan for mAbs from group **A, B, C, or D,** respectively, and indicated. Residues that engage hACE2 are highlighted in tan.

### Anti-SARS-CoV-2 mAbs protect against virus challenge *in vivo*

We next tested the anti-SARS-CoV-2 mAbs for protection against the historical SARS-CoV-2 WA1/2020 virus *in vivo*. Eight- to ten-week-old K18 human ACE2 (hACE2) transgenic mice were administered a single 100 µg dose (∼5 mg/kg) of anti-SARS-CoV-2 mAb 24 h prior to intranasal inoculation with 10^3^ FFU of SARS-CoV-2 WA1/2020. Mice treated with the isotype control mAb lost up to 25% body weight by 7 days post-infection (dpi), the designated endpoint of the study (**Fig 4A**). Mice treated with group A mAbs SARS2-71 and SARS2-34 maintained body weight until 6-7 dpi, at which point we observed a 10% weight loss (**Fig 4A**). Mice treated with group B mAbs SARS2-02 and SARS2-55 and group C mAbs SARS2-01 and SARS2-38 all maintained body weight throughout the experiment (**Fig 4B-C**). Animals treated with group D mAbs SARS2-10, SARS2-31 or SARS2-03 generally were less protected against virus-induced weight loss (**Fig 4D**).

**Figure 4.**
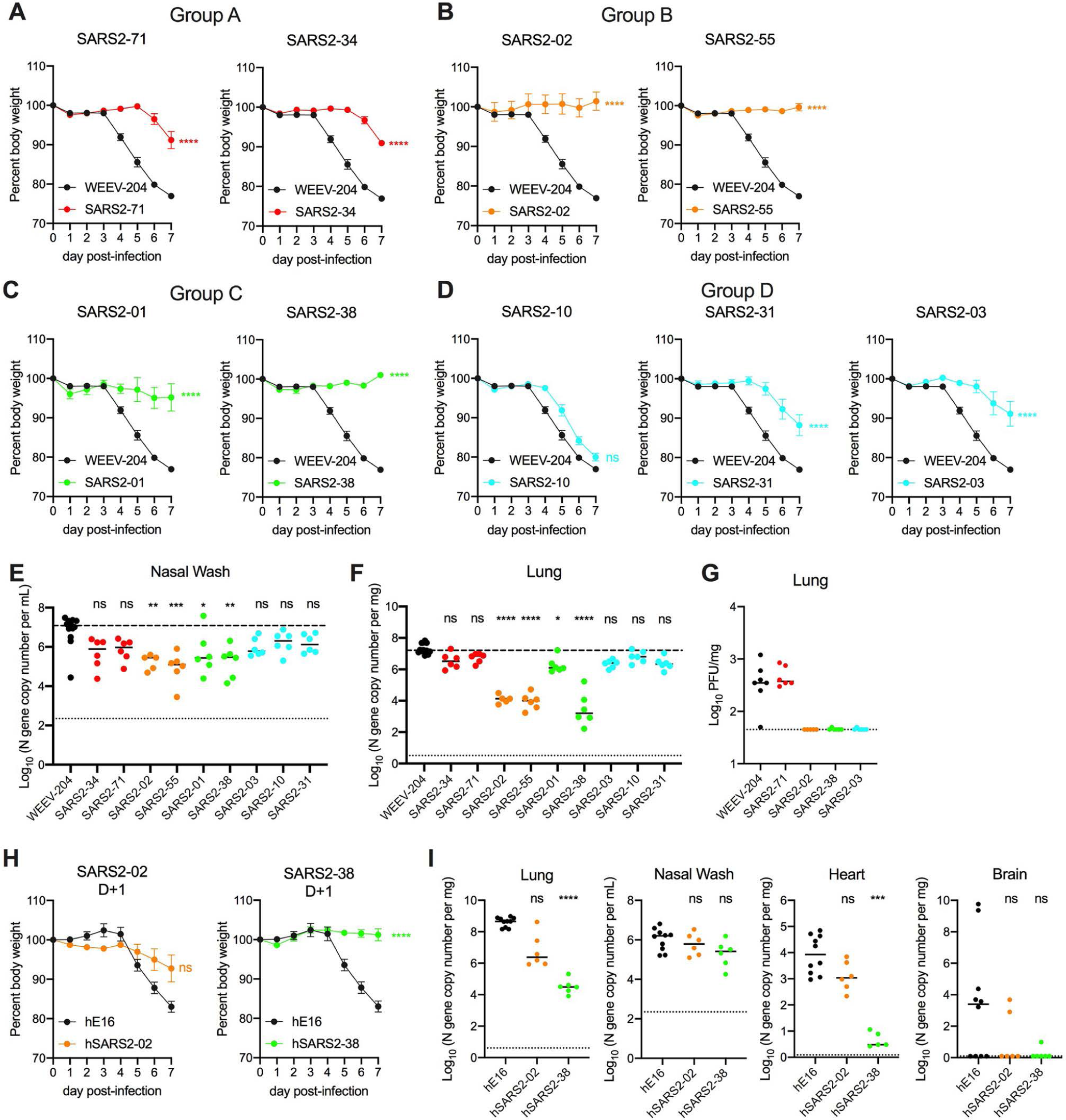
Anti- SARS-CoV-2 mAbs protect against SARS-CoV-2 infection *in vivo*. (**A-G**) K18-hACE2 transgenic mice were passively administered 100 µg (5 mg/kg) of the indicated mAb by intraperitoneal injection 24 h prior to intranasal inoculation with 10^3^ FFU of SARS-CoV-2 WA1/2020. (**A-D**) Mice were monitored for weight change for 7 days following viral infection. Mean weight change is shown. Error bars represent SEM. (**E-F**) At 7 dpi, nasal washes (**E**), and lungs (**F**) were collected, and viral RNA levels were determined. Median levels are shown; top dotted line indicates median viral load of control mAb-treated mice; bottom dotted line represents the limit of detection (LOD) of the assay. (**H**) A subset of the lungs from (**F**) were assessed for infectious viral burden by plaque assay. Median PFU/mL is shown. Dotted line indicates the LOD. (**A-F**) Data for each mAb are from two experiments; WEEV-204 (isotype control): n = 12; all other mAbs: n = 5-6 per group. (**H-I**) K18-hACE2 transgenic mice were passively given 200 µg (10 mg/kg) of the indicated mAb by intraperitoneal injection 24 h after intranasal inoculation with 10^3^ PFU of SARS-CoV-2 WA1/2020. Data are from two or three experiments; WEEV-204 (isotype control): n = 10; SARS2-02 and SARS2-38: n = 6 per group. (**H**) Mean weight change is shown. Error bars represent SEM. (**I**) At 7 dpi, lung, nasal washes, heart, and brain were collected and viral RNA levels were determined. (**A**-**D** and **H**) One-way ANOVA with Dunnett’s post-test of area under the curve. ns, not significant; ****p<0.0001. (**E, F, and I**) Kruskal-Wallis with Dunn’s post-test: ns, not significant, *p<0.05, ** p<0.01, ***p < 0.001, ****p < 0.0001.

To corroborate these findings, we measured the effect of mAb treatment on viral burden in the nasal washes and lungs on 7 dpi. The greatest decreases in viral RNA levels (∼30 to 100-fold) in the nasal washes relative to isotype control mAb-treated mice were observed in animals treated with mAbs in group B (SARS2-02 and SARS2-55) and group C (SARS2-01 and SARS2-38) (**Fig 4E**). The largest reductions in viral RNA levels in the lung (∼100 to 1,000-fold) again were observed for mice treated with mAbs in group B (SARS2-02 and SARS2-55) and group C (SARS2-38) (**Fig 4F**). A smaller (∼10-fold) decrement of virus RNA levels in the lung was observed for group C mAb SARS2-03. We also measured effects on infectious viral load in the lung by plaque assay for a subset of representative mAbs from each group. Whereas group A mAb SARS2-71 did not decrease the number of plaque-forming units (PFU) in the lung relative to the isotype control mAb-treated mice, SARS2-02, SARS2-38, and SARS2-03 all reduced infectious virus levels in the lung to the limit of detection of the assay (**Fig 4G**). The lack of protection conferred by SARS2-71 *in vivo* was unanticipated given its potent neutralizing activity in cell culture (EC50 of 8 ng/mL, **Fig 2**). Sequencing of viral RNA from the lungs of SARS2-71-treated mice at 7 dpi revealed an S477N mutation in the RBD in all samples, which was not present in the input WA1/2020 virus. Notably, S477N also emerged *in vitro* as an escape mutant under SARS2-71 selection pressure using the VSV-eGFP-SARS-CoV-2-S virus (**Fig 3A**). Thus, despite its potent inhibitory activity *in vitro*, SARS2-71 likely failed to protect *in vivo* because of rapid emergence of a fully pathogenic escape mutant.

To evaluate further the level of protection conferred by a subset of mAbs in our panel, we measured levels of cytokines and chemokines in lung tissues at 7 dpi, which are markers of the inflammatory and pathological outcomes in this mouse model (Golden et al., 2020; Oladunni et al., 2020; Winkler et al., 2020; Yinda et al., 2021). SARS2-38 and SARS2-02 treatment resulted in substantially reduced cytokine and chemokine levels relative to isotype control mAb-treated mice, with levels equivalent to those seen in naïve mice (**Fig S3**). In contrast, treatment with SARS2-71 and SARS2-03 did not result in these reductions, with cytokine and chemokine levels similar to isotype-control treated and infected mice.

To test for post-exposure therapeutic protection against SARS-CoV-2 challenge, we cloned the variable regions of group B mAb SARS2-02 and group C mAb SARS2-38 and inserted them into a human IgG1 backbone to make chimeric antibodies. We did this since chimeric, humanized, or fully-human mAbs are more likely to be used in humans, and because Fc effector functions contribute to the therapeutic activity of neutralizing SARS-CoV-2 mAbs *in vivo* (Winkler et al., 2021); the original murine IgG1 isotype of these mAbs binds poorly to the activating murine FcγRI and FcγRIV, whereas human IgG1 binds these murine Fc receptors with higher affinity and thus could have enhanced effector function (Dekkers et al., 2017). We confirmed the neutralizing activity of the chimeric mAbs hSARS2-02 and hSARS-38 relative to the original murine IgG1 versions of the mAbs (**Fig S4**). Next, we inoculated K18-hACE2 mice with 10^3^ FFU of SARS-CoV-2 WA1/2020. Twenty-four hours later, we administered a single 200 µg (10 mg/kg) dose of hSARS2-02, hSARS2-38, or an isotype control mAb. Both hSARS2-02 and hSARS2-38 protected against weight loss following infection (**Fig 4H**). At 7 dpi, hSARS2-38 reduced viral RNA levels in the lung and heart by ∼10,000-fold, whereas hSARS2-02 reduced infection by only ∼10-100 fold in these tissues (**Fig 4I**).

### Neutralization of variants of concern by anti-SARS-CoV-2 mAbs

We tested the two mAbs (SARS2-02 and SARS2-38) that conferred the greatest protection against WA1/2020 *in vivo* for neutralization of viruses with spike proteins corresponding to circulating variants of concern (VOCs). Recombinant chimeric WA1/2020 viruses encoding the spike protein from B.1.351 or B.1.1.28 were utilized for these studies (Wash-1.351 and Wash-1.1.28), as well as WA1/2020 with an introduced D614G mutation; we also tested viral isolates B.1.1.7, B.1.429, B.1.1.298, and B.1.222. Several of these VOCs encode amino acid changes in spike that can affect mAb binding (**Fig 5A**) (Chen *et al*., 2021d; Shen et al., 2021; Wang et al., 2021), including changes we identified in our VSV-eGFP-SARS-CoV-2-S escape mutant panel: L452R and E484K both showed reduced sensitivity to neutralization of VSV-eGFP-SARS-CoV-2-S by group B mAbs SARS2-02 and SARS2-55. Indeed, SARS2-02 exhibited reduced (∼50-100-fold) neutralizing activity against authentic SARS-CoV-2 strains with E484K (Wash-B.1.351 and Wash-B.1.1.28) or L452R (B.1.429) substitutions (**Fig 5B and C**). Notably, SARS2-38 did not lose potency against any of the variant viruses, with EC50 values ranging from 1-4 ng/mL across the panel tested (**Fig 5D and E**).

**Figure 5.**
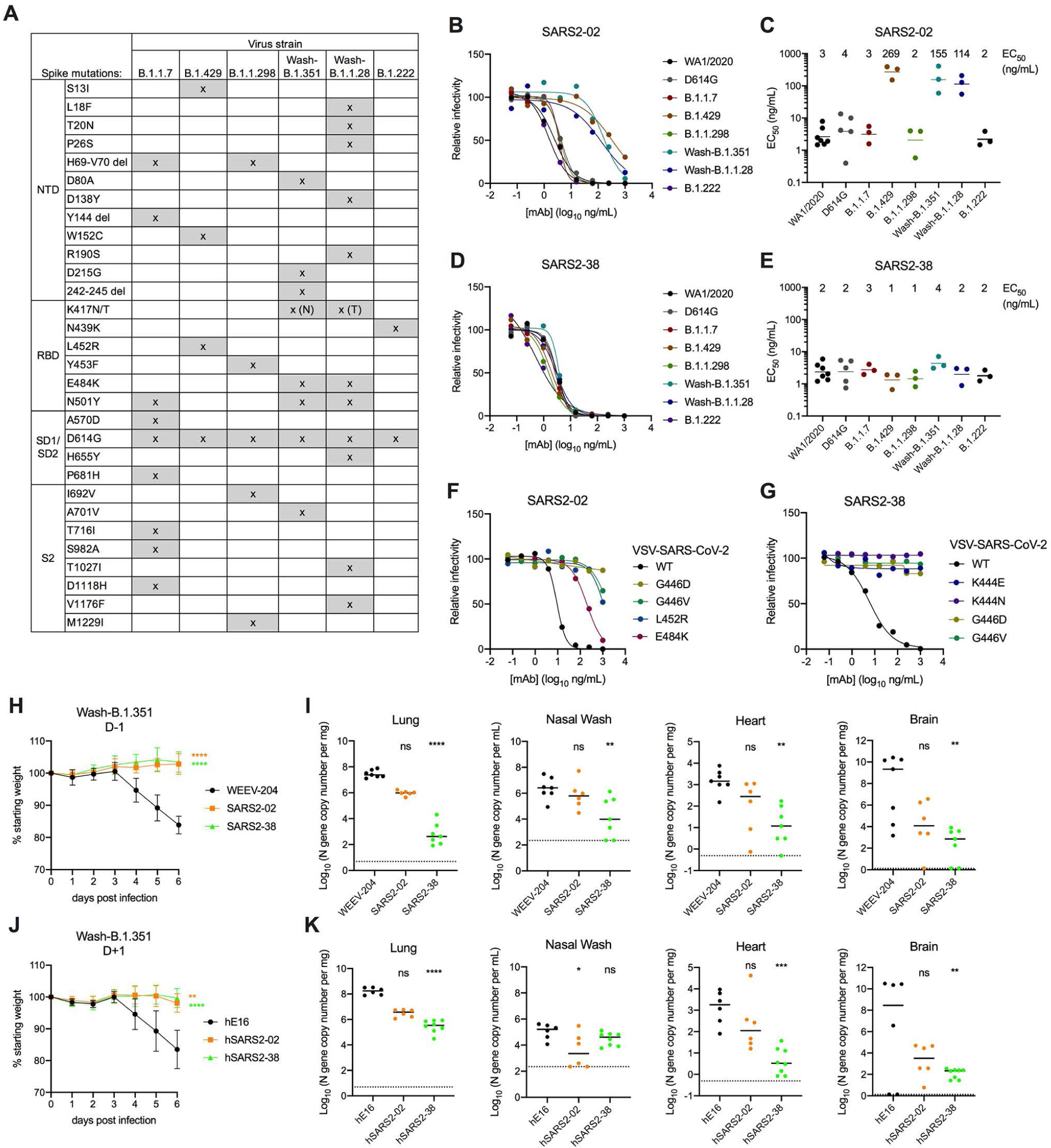
Neutralization of variants of concern by anti-SARS-CoV-2 mAbs. (**A**) Variants of concern (VOC) and their mutations in spike. SARS2-2 (**B-C**) and SARS2-38 (**D-E**) were tested for neutralization of the indicated variants by FRNT. (**B and D**) Representative dose response curves are shown. (**C and E**) Mean EC50 values are shown; data are from three to five experiments. (**F-G**) Representative dose response curves of SARS2-02 and SARS2-38 neutralization of VSV-eGFP-SARS-CoV-2-S and the indicated neutralization-resistant mutants. Data is from of one of two experiments. (**H-I**) K18-hACE2 mice were administered 100 µg (5 mg/kg) of the indicated mAb by intraperitoneal injection 24 h prior to intranasal inoculation with 10^3^ FFU of SARS-CoV-2 Wash-B.1.351. (**H**) Mean weight change is shown. Error bars represent SEM. (**I**) At 6 dpi, the indicated tissues were collected, and viral RNA levels were determined. Data are from two experiments; WEEV-204 (isotype control) and SARS2-38: n = 7; SARS2-02: n = 6. (**J-K**) K18-hACE2 mice were inoculated with 10^3^ FFU of SARS-CoV-2 Wash-B.1.351, and 24 h later they were administered 200 µg (10 mg/kg) of the indicated mAb. (**J**) Mean weight change is shown. Error bars represent SEM. (**K**) At 6 dpi, the indicated tissues were collected, and viral RNA levels were determined. Data are from two experiments; hWNV-E16 and hSARS2-02: n=6; hSARS2-38: n=8). (**H** and **J**) One-way ANOVA with Dunnett’s post-test of area under the curve. **p<0.01; ****p<0.0001. (**I and K**) Kruskal-Wallis with Dunn’s post-test: ns, not significant, *p < 0.05, **p < 0.01, ***p<0.001, ****p < 0.0001.

To expand on this analysis, we tested the VSV-eGFP-SARS-CoV-2-S viruses that were resistant to SARS2-02 and SARS2-38 for neutralization using full dose response curves analysis. SARS2-02 showed ∼20-fold reduced potency against E484K, ∼100-fold reduced potency against L452R and G446V, and did not neutralize G446D at the highest concentration of mAb tested (**Fig 5F**). SARS2-38 showed virtually no neutralizing activity against K444E, K444N, G446D, or G446V mutants even at the highest concentration (1 μg/ml) of mAb tested (**Fig 5G**). Despite these results with VSV-eGFP-SARS-CoV-2-S viruses, when we serially passaged authentic WA1/2020 D614G or Wash-B.1.351 SARS-CoV-2 in Vero-TMPRSS2-ACE2 cells in the presence of neutralizing mAbs, we readily isolated resistant viruses following SARS-02 but not SARS2-38 selection with both strains.

We tested SARS2-02 and SARS2-38 for protection against Wash-B.1.351 in K18-hACE2 mice. Animals treated with 100 µg of either SARS2-02 or SARS2-38 24 h prior to infection were protected from weight loss (**Fig 5H**), despite the reduced neutralization potency of SARS2-02 against Wash B.1.351. SARS2-38 treatment greatly reduced viral titers in the lung, nasal washes, heart, and brain at 7 dpi compared to the isotype control-treated mice, whereas SARS2-02 had less of a protective effect (**Fig 5I**). When hSARS2-02 and hSARS2-38 were administered to the K18-hACE2 transgenic mice as therapy 24 h after infection with Wash-B.1.351, a similar phenotype was observed: while they both protected mice against weight loss (**Fig 5J**), hSARS2-38 resulted in a greater reduction in viral titers at 7 dpi in the lung, heart, and brain than hSARS2-02 (**Fig 5K**).

### SARS2-38 targets the proximal RBM ridge with extensive light chain contact

To define further the mechanistic basis for the broad and potent neutralization by SARS2-38, we first analyzed the interaction of antigen binding fragments (Fab) of SARS2-38 with SARS-CoV-2 spike using biolayer interferometry (BLI). SARS2-38 bound spike with high monovalent affinity (kinetically derived K_D_ of 6.5 nM) and had a half-life of 4.8 min (**Fig S5A**). To understand the basis for this binding structurally, we performed cryo-electron microscopy (cryo-EM) on complexes of SARS2-38 Fab and the SARS-CoV-2 spike protein (**Fig S5B**). We generated three-dimensional classes to sample the conformational landscape of the Fab/spike complex, and the class of highest resolution was refined further. This class consisted of trimeric spike with all RBDs in the down position (D/D/D) and one RBD bound by Fab (**Fig 6A and S6A-B**). Using non-uniform refinement, we achieved an overall resolution of 3.20 Å, with local resolution ranging from ∼2.5 Å in the core of the spike to ∼5.5 Å in the constant region of the Fab, which was visible only at high contour (**Fig S6B-D**). Other binding configurations also were seen, the most predominant consisting of spike with one RBD up and two RBDs down (U/D/D), with only the up RBD bound by Fab (31.1%). Less frequently, all three RBDs were bound by Fab in the U/D/D conformation (22.0%; **Fig S5B**). Although SARS2-38 could bind SARS-CoV-2 spike with full occupancy, 61.1% of spike trimers were bound only by a single Fab molecule.

**Figure 6.**
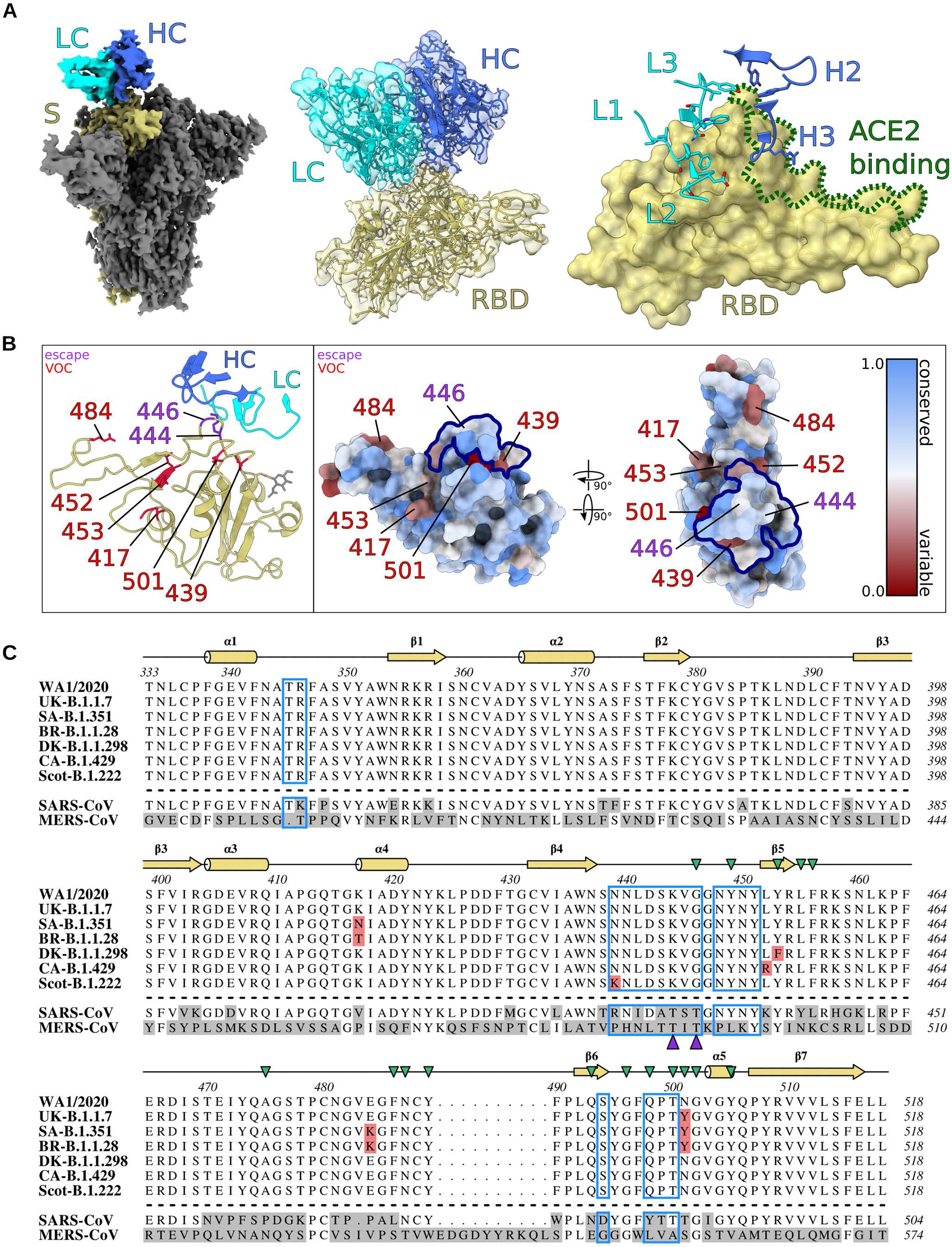
SARS2-38 targets a conserved portion of the RBM with extensive light chain contact. (**A**) *Left panel*. Density map of SARS2-38 Fv bound to trimeric SARS-CoV-2 spike protein with all RBDs in the down position. The spike monomer bound by SARS2-38 is shown in yellow with the rest of the trimer colored gray. The SARS2-38 heavy chain is shown in royal blue, and the light chain in cyan. *Middle panel*. Focused density map of the Fv/RBD complex encompassing a refined atomic model. The RBD is shown in yellow. The SARS2-38 heavy and light chains are colored royal blue and cyan, respectively. *Right panel*. Complementarity-determining regions (CDRs) of SARS2-38 overlay a surface rendering of the RBD. CDRs from the heavy and light chains are colored royal blue and cyan, respectively, with the RBD colored yellow. ACE2-binding residues of the receptor binding motif (RBM) are outlined in green. (**B**) *Left panel*: a ribbon diagram of the RBD and SARS2-38 CDRs with escape mutations and variants of concern (VOCs) noted in purple and red, respectively. The RBD is otherwise colored yellow, with a gray glycan linked to N343. CDRs of the SARS2-38 heavy and light chain are colored royal blue and cyan, respectively. *Right panel*: surface renderings of RBD colored according to conservation of surface residues (blue = conserved, red = variable). Escape mutations and VOCs are noted in purple and red, respectively. The SARS2-38 epitope is outlined in navy. (**C**) Multiple sequence alignment of RBD (residues 333-518) from WA1/2020, SARS-CoV-2 VOCs, SARS-CoV, and MERS-CoV, with the binding footprint of SARS2-38 boxed in blue. Mutations within SARS-CoV-2 VOCs are highlighted in red, and SARS2-38 escape mutation contacts are marked with purple triangles. Secondary structure annotation is displayed above the alignment in yellow with ACE2 contacts designated by green triangles (Lan et al., 2020). Divergent residues within SARS-CoV and MERS-CoV (relative to WA1/2020) are highlighted in gray.

To improve resolution at the Fab/RBD interface in the D/D/D reconstruction, we performed a focused, local refinement of the SARS2-38 variable domain (Fv) and RBD, excluding the rest of the spike and the constant region of the Fab. This reconstruction of the Fv/RBD complex achieved a resolution of 3.16 Å, allowing unambiguous placement of the protein backbone, secondary structures, and most side chains at the interface (**Fig S5B and S6E-F**). The SARS2-38 Fv sits atop three loops protruding at the proximal end of the RBM between helix α1 and strand β1 (contact residues T345-R346), strands β4 and β5 (N439-G446, N448-Y451), and strand β6 and helix α5 (S494 and Q498-T500; **Fig 6A-B**); these results correspond well with our VSV-based escape mutant mapping (**Fig 3**). All three light chain CDRs contact loop β4-β5, with CDR2 and CDR3 forming additional contacts with loops α1-β1 and β6-α5, respectively. In comparison, the heavy chain interacts in a more limited manner with loops β4-β5 and β6-α5 via CDR2 and CDR3. CDR1 of the heavy chain makes no contact at all with the RBD. The heavy chain does, however, engage ACE2 contact residues of the RBM **(Fig 6A**, *right panel*). This and other steric effects likely explain the inhibition of ACE2 binding by SARS2-38.

### The SARS2-38 epitope is conserved among circulating SARS-CoV-2 variants of concern

SARS2-38 potently neutralized all tested VOCs. To understand this broadly-neutralizing activity, we mapped the SARS2-38 epitope alongside VOC mutations within the RBD (**Fig 6B**, *left panel* **and Fig 6C**). One mutation in the SARS2-38 footprint, N439K, is present in variant B.1.222 and resides at the periphery of the epitope. However, B.1.222 remained sensitive to neutralization by SARS2-38, and escape mutants at this residue were not generated *in vitro*, suggesting that N439 is not critical for SARS2-38 binding. The SARS2-38 epitope includes no other residues corresponding to VOC mutations, which explains its performance against these variants. Notwithstanding this, we could select escape mutations *in vitro* in the context of VSV-eGFP-SARS-CoV-2-S chimeric virus, namely K444E/N and G446D/V substitutions, which reside on the β4-β5 loop central to the SARS2-38 epitope (**Fig 6B-C**). The substitutions generated at K444 result in a loss of positive charge (K444N) or charge reversal (K444E), whereas the mutation at G446 may distort the entire loop structure; in our model, G446 adopts a stereochemistry unique to the glycine residue (φ= 108° and =Ψ-19°). This structural analysis likely explains the resistance conferred by these amino acid substitutions.

To understand the efficacy of SARS2-38 amidst the landscape of all circulating variants, we used the COVID-19 CoV Genetics Browser (covidcg.org) to probe RBD sequences in the GISAID database (786,273 isolates as of March 28, 2021; Chen et al., 2021; Shu et al., 2017). We then developed a log-scale conservation score for RBD residues 333-520. In this model, perfect conservation of the reference amino acid (from 2019n-CoV/WA1/2020) across all isolates corresponds to a score of 1, and complete loss of the reference amino acid results in a score of 0. Visualizing these scores on a color-coded RBD surface rendering (blue = 1, more conserved; red = 0, more variable) revealed that the RBM is generally more variable than the rest of the RBD, with VOCs clearly seen as red patches (**Fig 6B**, *right panel*). This analysis also suggested that in addition to not being affected by the VOCs tested in this study, SARS2-38 targets a portion of the RBM that is conserved among circulating SARS-CoV-2 variants. The positions at which we identified escape mutants using VSV-eGFP-SARS-CoV-2-S chimeric viruses were substituted in only 0.02% (K444) and 0.04% (G444) of isolates, with the specific escape mutations (K444E/N and G446D/V) observed in only 0.007% and 0.03% of isolates respectively. Overall, 99.96% of isolates lacked the escape mutations for SARS2-38 identified in our study.

## DISCUSSION

In this study, we describe and characterize extensively a panel of mAbs that bind the RBD of the SARS-CoV-2 spike protein. Several anti-RBD mAbs protected *in vivo* against SARS-CoV-2 infection in K18-hACE2 transgenic mice. While the less potently neutralizing mAbs directed against epitopes on the base of RBD (SARS2-10, SARS2-31, and SARS2-03) exhibited diminished protection against weight loss, induction of inflammatory cytokines and chemokines in the lung, and viral infection in the lung and nasal wash than mAbs recognizing the RBM, neutralization potency was not the only predictor of *in vivo* efficacy. Indeed, SARS2-71 neutralized SARS-CoV-2 with a potency similar to that of protective mAbs SARS2-02 and SARS2-38, yet failed to confer protection in mice. Notwithstanding this result, antibodies targeting proximal competing epitopes as SARS2-71, such as COV2-2196, have been shown to confer protection *in vivo* (Zost *et al*., 2020). The failure of SARS2-71 to protect in particular is likely due to the emergence of the escape variant S477N *in vivo*. This finding demonstrates that SARS-CoV-2 can rapidly escape from mAb inhibition *in vivo*, and that mAb or mAb cocktails that prevent or limit rapid escape mutant generation likely will have greater therapeutic utility. While currently authorized mAb treatments include cocktails, the emergence of VOCs that are resistant to one or both component mAbs could compromise drug efficacy.

The most potently inhibitory mAbs in our panel bind epitopes within or proximal to the RBM and inhibit spike interaction with human ACE2 by ELISA, as observed for other anti-SARS-CoV-2 mAbs (Zost *et al*., 2020). Several of these mAbs inhibited viral attachment to Calu-3 and Vero-TMPRSS2-ACE2 cells, but not to Vero E6 cells or Vero-TMPRSS2 cells. Infection of Vero E6 cells by SARS-CoV-2 is dependent on endogenous levels of monkey ACE2 expression, as pretreatment with anti-ACE2 mAbs inhibits infection (Hoffmann et al., 2020). However, other host factors such as heparan sulfate also can mediate virus attachment to cells (Chu et al., 2021; Clausen et al., 2020). If binding to other cell surface ligands occurs prior to the RBD-ACE2 interaction, mAbs that block ACE2 binding may not efficiently inhibit SARS-CoV-2 attachment, but instead block a downstream ACE2-dependent entry step. This idea is supported by our data showing that several neutralizing mAbs block viral internalization in Vero E6 cells. Moreover, anti-RBD mAbs have only moderate decreases in neutralization potency when added after virus absorption to Vero E6 cells. In contrast, when SARS-CoV-2 attaches to the cell surface via human ACE2 interaction, such as in Vero-TMPRSS2-ACE2 cells, the addition of anti-RBD mAbs after attachment failed to neutralize virus infection. A higher density of ACE2 or higher affinity of spike protein for human ACE2 (relative to monkey ACE2) on the Vero-TMPRSS2-ACE2 cells may drive initial virus attachment through the RBD-ACE2 interaction and explain why mAbs can block this step in these cells. Together, these data suggest that the ability of anti-RBD mAbs to inhibit SARS-CoV-2 attachment depends on cellular ACE2 expression levels and thus can be cell-type dependent. As these mechanistic differences did not markedly affect mAb potency on the different cellular substrates, we conclude that in the cells we tested there is a required entry interaction with ACE2 either at attachment, post-attachment, or internalization steps.

Several mutations and deletions in emerging VOCs occur in the NTD and RBD that allow them to avoid antibody recognition, including RBD mutations K417N/T (B.1.351 and B.1.1.28), N439K (B.1.222), L452R (B.1.429), Y453R (B.1.1.298), E484K (B.1.351 and B.1.1.28), and N501Y (B.1.1.7, B.1.351, and B.1.1.28) (reviewed by (Plante et al., 2021)), highlighting the importance of developing mAbs against a variety of spatially distinct epitopes. In our panel, SARS2-38 potently neutralized viruses encoding any of the above mutations, did not readily select for escape mutations with authentic SARS-CoV-2 strains, and retained therapeutic activity *in vivo* against a virus containing substitutions of one of the key VOCs (B.1.351). Moreover, functional mapping and structural analysis of the binding footprint of SARS-CoV-2 defined a conserved RBD epitope that could be recognized by other potently neutralizing and protective human mAbs.

Relatively few antibodies targeting similar epitopes to SARS2-38 have been described, and those characterized bind the RBD in distinct orientations with heavy chain predominance (**Fig 7A**). These include murine mAb 2H04, as well as human mAbs REGN10987, COV2-2130, and, though less similar, S309 (Dong *et al*., 2021; Hansen *et al*., 2020; Liu *et al*., 2021; Pinto et al., 2020). SARS2-38 differs in two respects: (a) the baseline neutralizing activity of SARS2-38 against WA1/2020 in Vero cells (EC50, ∼5 ng/mL) is 30-fold, 20-fold, and 16-fold more potent than that of 2H04, COV2-2130, and S309, respectively (Alsoussi *et al*., 2020; Pinto *et al*., 2020; Zost *et al*., 2020); and (b) SARS2-38 retains strong neutralization potency against all VOCs evaluated in this study, whereas the inhibitory activity 2H04, COV2-2130, and S309 is reduced somewhat against B.1.1.7, B.1.429, and B.1.351, respectively ((Chen et al., 2021c; Chen *et al*., 2021d) and R.E.C. and M.S.D. unpublished results). Similarly, REGN10987 exhibited a 10-fold reduction in neutralizing activity against B.1.429 compared to WA1/2020 (Chen *et al*., 2021c; Hansen *et al*., 2020; Wang *et al*., 2021). A structural examination of these other antibody footprints within the context of VOC mutations does not provide a direct explanation for some of the resistance (**Fig 7B**). Instead, allostery may play a role. Whereas other broadly and potently neutralizing mAbs (including mAbs 2C08, COV2-2196, 58G6, 510A5, and S2X259) have been reported that bind RBD epitopes at residues G476, F486, and N487, or loops near residues 369-386, 404-411, 450-458, and 499-508 (Dong *et al*., 2021; Li et al., 2021; Schmitz et al., 2021; Tortorici et al., 2021), SARS2-38 targets a distinct epitope proximal to the RBM and has been evaluated functionally against a larger panel of authentic viruses containing sequences corresponding to emerging SARS-CoV-2 variants.

**Figure 7.**
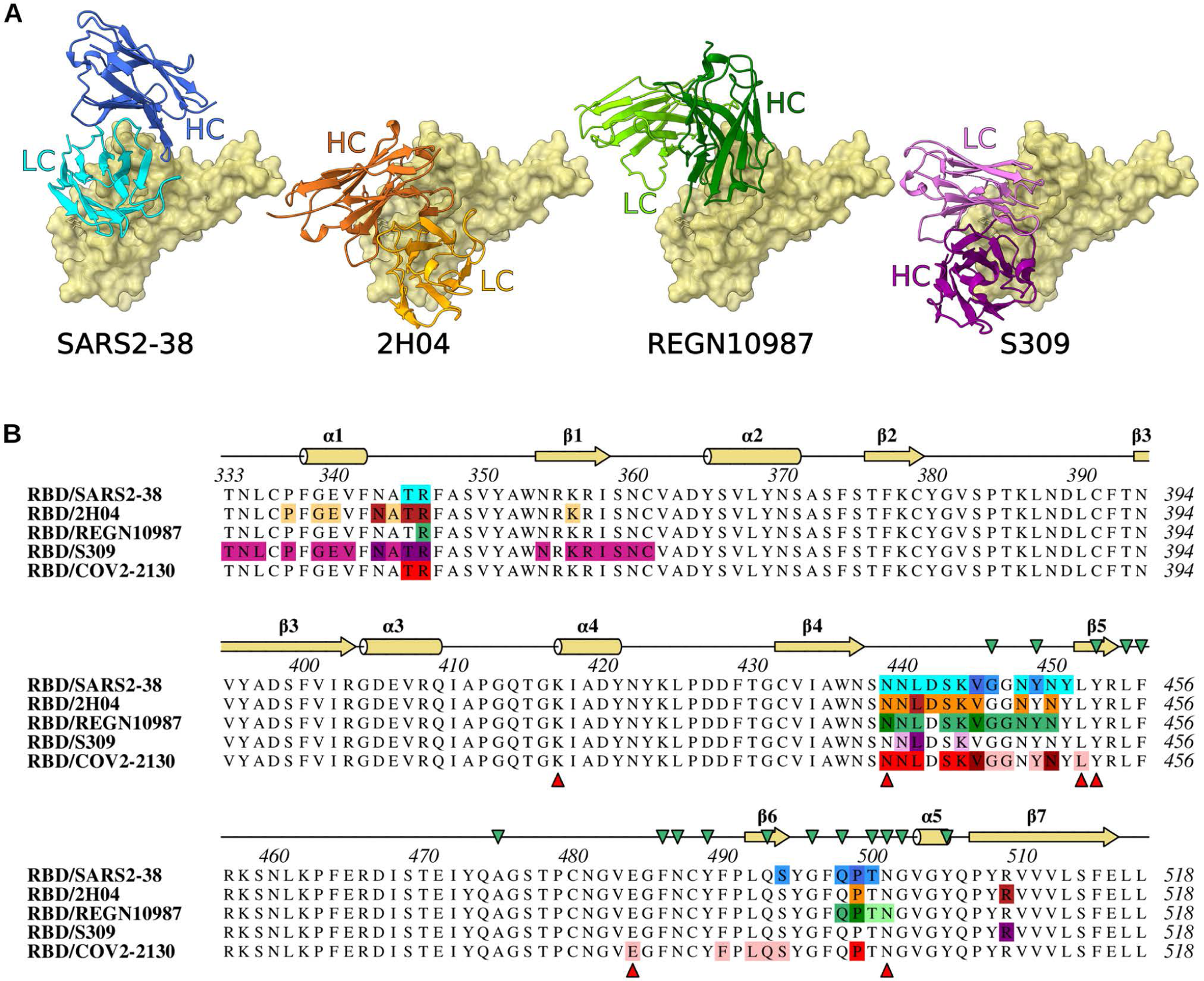
Similarity of SARS2-38 epitope to other mAbs. (**A**) Structural comparison of SARS2-38 to mAbs targeting a similar region of the RBD. (**B**) Multiple sequence alignment of the SARS-CoV-2 RBD (residues 333-518) with mAb binding footprints as determined by qtPISA analysis. For SARS2-38, heavy chain, light chain, and shared contacts are shown in blue, cyan, and dark blue, respectively. For 2H04, heavy chain, light chain, and shared contacts are shown in orange, pale orange, and dark orange, respectively. For REGN10987, heavy chain, light chain, and shared contacts are shown in green, pale green, and dark green, respectively. For S309, heavy chain, light chain, and shared contacts are shown in magenta, pale purple, and purple, respectively. For COV2-2130, heavy chain, light chain, and shared contacts are shown in red, pale red, and brick red, respectively. Secondary structure annotation is displayed above the alignment in yellow, with ACE2 contacts designated by green triangles (Lan et al., 2020). VOC substitutions are designated below the alignment by red triangles.

In summary, we have characterized a panel of anti-SARS-CoV-2 mAbs, defined their cellular mechanism of action in different cells, tested *in vitro* neutralizing and *in vivo* protection capacity against historical and circulating variants, and determined the structure of the viral spike protein bound to SARS2-38, a potently and broadly neutralizing mAb that recognizes emerging VOCs. A humanized version of SARS2-38 confers therapeutic protection against the WA1/2020 isolate and a SARS-CoV-2 strain expressing the spike protein of B.1.351. The conserved epitope bound by SARS2-38 thus may be a potential target for antibodies with therapeutic potential or that are induced by effective vaccines with more limited potential for resistance against VOCs.

## ACKNOWLEDGEMENTS

This study was supported by contracts and grants from NIH (75N93019C00062, HHSN272201700060C, 75N93019C00074, U01 AI151810, R01 AI118938, and R01 AI157155), and the Defense Advanced Research Project Agency (HR001117S0019). J.B.C. is supported by a Helen Hay Whitney Foundation postdoctoral fellowship. This work also was funded by a generous gift from Washington University. We gratefully acknowledge the originating and submitting laboratories who generated and shared genetic sequence data via the GISAID Initiative. We also thank Charles Chiu and Raul Andino for providing the B.1.429 isolate and Barney Graham for cell lines and experimental advice.

## AUTHOR CONTRIBUTIONS

Conceptualization and Methodology, L.A.V., M.S.D., L.J.A., D.H.F., Z.L., and S.P.J.W; Investigation, L.A.V., L.J.A., Z.L., R.E.C., P.G., S.R., B.K.S., J.B.C., B.M.W., L.D., and I.D.A.; Formal analysis, L.J.A. and S.A.H.; Key reagents, E.S.W., H.Z., P-Y.S., A.P., and A.C.; Supervision and funding, M.S.D., D.H.F., D.W., A.C.M.B., J.E.C., and S.P.J.W; Writing – original draft, L.A.V., L.J.A., and M.S.D.; Writing – review and editing, all authors.

## SUPPLEMENTAL FIGURE LEGENDS

**Figure S1. Competition profile of mAb panel, Related to Figure 1.** A subset of mAbs from each reference mAb competition group were tested for competition for SARS-CoV-2 spike binding against each other. Data represent mean of technical duplicates.

**Figure S2. Neutralization by anti-SARS-CoV-2 mAbs on different cell substrates, Related to Figure 2.** Anti-SARS-CoV-2 mAbs were assayed for neutralization by FRNT against SARS-CoV-2 using Vero E6, Vero-TMPRSS2, Vero-TMPRSS2-ACE2. (**A**) Representative dose response curves are shown. (**B**) Mean EC50 values are shown; data are from three experiments.

**Figure S3. Cytokine and chemokine levels in the lungs of SARS-CoV-2 infected mice following treatment with anti-SARS-CoV2 mAbs, Related to Figure 4.** Cytokine and chemokine levels in lung homogenates harvested in **Fig 5** were measured by a multiplex platform. (**A**) Heat map showing Log_2_ fold change in cytokine and chemokine levels compared to lungs from mock-infected animals. (**B**) Levels of each cytokine and chemokine are plotted. Data are from two experiments, n = 5-6 per group. One-way ANOVA with Dunnett’s post-test: ns, not significant, ∗p < 0.05, ∗∗p < 0.01, ∗∗∗p < 0.001, ∗∗∗∗p < 0.0001.)

**Figure S4. Validation of chimeric mAb activity, Related to Figure 4.** Anti-SARS-CoV-2 chimeric mouse Fv/human IgG1 Fc mAbs were assayed for neutralization by FRNT against SARS-CoV-2 WA/2020. m, mouse hybridoma-derived mAb, and h, recombinant chimeric mAb. Representative dose response curves are shown. Data are from three experiments.

**Figure S5. Binding analysis and cryo-EM data processing pipeline, Related to Figure 6.** (**A**) Biolayer interferometry signal (left) and steady state analysis (right) of SARS2-38 Fab interacting with immobilized SARS-CoV-2 spike. Kinetic values were fitted to a 1:1 binding model with a drifting baseline. (**B**) Flowchart depicting data processing steps for global reconstruction of SARS2-38 Fv bound to trimeric spike and local refinement of SARS2-38 Fv bound to RBD.

**Figure S6. Validation of global and local cryo-EM reconstructions of SARS2-38 Fv bound to SARS-CoV-2 spike/RBD, Related to Figure 6.** (**A**) Density map and fitted model of SARS2-38 Fv bound to trimeric SARS-CoV-2 spike. The spike monomer bound by SARS2-38 is shown in yellow, with the rest of the trimer colored gray. The SARS2-38 heavy chain is shown in royal blue, and the light chain in cyan. (**B**) Orientational distribution assigned to particles in global refinement of SARS2-38 Fv bound to trimeric spike. (**C**) GSFSC curve for global refinement of SARS2-Fv bound to trimeric spike. (**D**) Local resolution map for global refinement of SARS2-38 Fv bound to trimeric spike. (**E**) GSFSC curve for local refinement of SARS2-Fv bound to RBD. (**F**) Example density and model fits for an RBD beta strand (left) and at the SARS2-38/RBD interface (right). RBD is depicted in yellow, and the SARS2-38 light chain is shown in cyan.

**Table S1. Cryo-EM data collection, processing, and model refinement statistics, Related to Figure 6**. Respective statistics are provided for local and global refinements of SARS2-38 Fv bound to SARS-CoV-2 RBD/spike.

## STAR METHODS

### RESOURCE AVAILABLITY

**Lead Contact**. Further information and requests for resources and reagents should be directed to the Lead Contact, Michael S. Diamond.

**Materials Availability**. All requests for resources and reagents should be directed to the Lead Contact author. This includes mice, antibodies, viruses, and proteins. All reagents will be made available on request after completion of a Materials Transfer Agreement.

**Data and code availability.** All data supporting the findings of this study are available within the paper and are available from the corresponding author upon request. Structural datasets have been uploaded and are available at PDB (accession codes 7MKL and 7MKM).

## EXPERIMENTAL MODEL AND SUBJECT DETAILS

### Viruses

The 2019n-CoV/USA_WA1/2020 (WA1/2020) isolate of SARS-CoV-2 was obtained from the US Centers for Disease Control (CDC). WA1/2020 stocks were propagated on Vero CCL81 cells and used at passage 6 and 7. Viral titer was determined by focus-forming assay (FFA) on Vero E6 cells as described (Case *et al*., 2020). The D614G virus was produced by introducing the mutation into an infectious clone of WA1/2020, and the B.1.351 and B.1.1.28 Spike genes were cloned into the WA1/2020 infectious clone to produce Wash-B.1.351 and Wash-B.1.1.28 chimeric viruses, as described previously (Chen *et al*., 2021d). The B.1.1.7, B.1.429, B.1.298, and B.1.222 isolates were isolated from infected individuals. Viruses were propagated on Vero-TMPRSS2 cells and subjected to deep sequencing to confirm the presence of the substitutions indicated in **Fig 5A**.

### Cells

Cell lines were maintained at 37°C in the presence of 5% CO2. Vero E6 cells were passaged in Dulbecco’s Modified Eagle Medium (DMEM) (Invitrogen) supplemented with 10% fetal bovine serum (FBS) (Omega Scientific) and 100 U/mL penicillin-streptomycin (P/S) (Invitrogen). Vero cells that over-express TMPRSS2 or TMPRSS2-ACE2 were maintained as Vero CCL81 cells, with the addition of 5 µg/mL blasticidin (Vero-TMPRSS2) or 10 µg/mL puromycin (Vero-TMPRSS2-ACE2). Calu-3 cells were maintained in DMEM with 20% FBS and 100 U/mL P/S.

### Proteins

Genes encoding SARS-CoV-2 spike protein (residues 1-1213, GenBank: MN908947.3) and RBD (residues 319-514) were cloned into a pCAGGS mammalian expression vector with a C-terminal hexahistidine tag. The spike protein was prefusion stabilized and expression optimized via six proline substitutions (F817P, A892P, A899P, A942P, K986P, V987P) (Hsieh et al., 2020), with a disrupted S1/S2 furin cleavage site and a C-terminal foldon trimerization motif (YIPEAPRDGQAYVRKDGEWVLLSTFL). Expi293F cells were transiently transfected, and proteins were recovered via cobalt-charged resin chromatography (G-Biosciences) as previously described (Alsoussi et al., 2020; Hassan et al., 2020). For ACE2 binding inhibition analysis, the SARS-CoV-2 spike protein was made by synthesizing a gene encoding the ectodomain of a prefusion conformation-stabilized SARS-CoV-2 spike (S6Pecto) protein (Hsieh *et al*., 2020) containing C-terminal Twin-Strep-tag. The spike gene was then cloned it into a DNA plasmid expression vector for mammalian cells. Protein was produced in FreeStyle 293-F cells (Thermo Fisher Scientific) and purified from culture supernatants using StrepTrap HP affinity column (Cytiva).

### Mice

Animal studies were carried out in accordance with the recommendations in the Guide for the Care and Use of Laboratory Animals of the National Institutes of Health. The protocols were approved by the Institutional Animal Care and Use Committee at the Washington University School of Medicine (Assurance number A3381-01). Virus inoculations were performed under anesthesia that was induced and maintained with ketamine hydrochloride and xylazine, and all efforts were made to minimize animal suffering.

K18-hACE2 transgenic mice were purchased from Jackson Laboratories (#034860) and housed in a pathogen-free animal facility at Washington University in St. Louis. For passive transfer studies, mAbs were diluted in PBS and administered to mice via intraperitoneal injection in a 100 µL total volume. Viral infections were performed via intranasal inoculation with 10^3^ FFU of virus. Mice were monitored daily for weight loss.

## METHOD DETAILS

### MAb generation

BALB/c mice were immunized with 10 µg of SARS-CoV-2 RBD adjuvanted with 50% AddaVax™ (InvivoGen), via intramuscular route (i.m.), followed by i.m. immunization two and four weeks later with SARS-CoV-2 spike protein (5 µg and 10 µg, respectively) supplemented with AddaVax™. Mice received a final, non-adjuvanted boost of 25 µg of SARS-CoV-2 spike or RBD (12.5 µg intravenously and 12.5 µg interperitoneally) 3 days prior to fusion of splenocytes with P3X63.Ag.6.5.3 myeloma cells. Hybridomas producing antibodies that bound to SARS-CoV-2-infected permeabilized Vero CCL81 cells by flow cytometry and to SARS-CoV-2 recombinant spike protein by direct ELISA were cloned by limiting dilution. All hybridomas were screened initially with a single-endpoint neutralization assay using hybridoma supernatant diluted 1:3 and incubated with SARS-CoV-2 for 1 h at 37°C prior to addition to Vero E6 cells. Following a 30-h incubation, cells were fixed, permeabilized, and stained for SARS-CoV-2 infection with CR3022 as described (Case *et al*., 2020). A subset of neutralizing hybridoma supernatants were purified commercially (Bio-X Cell) after adaptation for growth under serum-free conditions.

### VSV-eGFP-SARS-CoV-2-S escape mutants

VSV-eGFP-SARS-CoV-2-S escape mutants were produced as described previously (Liu *et al*., 2021). Briefly, plaque assays were performed to isolate escape mutants on Vero-TMPRSS2 cells with neutralizing mAb in the overlay. Escape clones were plaque-purified on Vero-TMPRSS2 cells in the presence of mAb. Plaques in agarose plugs and viral stocks were amplified on MA104 cells at an MOI of 0.01 in Medium 199 containing 2% FBS and 20 mM HEPES pH 7.7 (Millipore Sigma) at 34°C. Viral supernatants were harvested upon extensive cytopathic effect and clarified of cell debris by centrifugation at 1,000 x g for 5 min.

### Determination of mAb concentration in hybridoma supernatant

The mAb concentration in each hybridoma supernatant was quantified by ELISA. Nunc MaxiSorp plates (Thermo Fisher Scientific) were coated with 1μg/mL of goat anti-mouse IgG (Southern Biotech) in 50 μL of NaHCO_3_ (pH 9.6) coating buffer and incubated overnight at 4°C. Plates were washed three times with ELISA wash buffer (PBS containing 0.05% Tween-20), and then incubated with 200 μL of blocking buffer (PBS, 2% BSA, 0.05% Tween-20) for 1 h at room temperature. Plates were incubated with hybridoma supernatant diluted 1:500 or 1:2000 in blocking buffer, or serial dilutions of purified isotype control mAb as a standard, for 1 h at room temperature. Plates were washed three times with ELISA wash buffer, and incubated with 50 μL of anti-mouse IgG-HRP (Sigma) diluted 1:500 for 1 h at room temperature. Plates were washed three times with ELISA wash buffer and three times with PBS, before incubation with 100 μL of TMB substrate (Thermo Fisher Scientific) for 3 min at room temperature before quenching with the addition of 50 μL of 2 N H_2_SO_4_ and measuring OD 450 nm. Antibody concentrations in hybridoma supernatant were interpolated from a standard curve produced using an isotype control mAb.

### Spike and RBD binding analysis

96-well Maxisorp plates were coated with 2 µg/mL of SARS-CoV-2 spike or RBD protein in 50 mM Na_2_CO_3_ (70 μL) overnight at 4 °C. Plates were washed three times with PBS + 0.05% Tween-20 and blocked with 200 μL of PBS + 0.05% Tween-20 + 1% BSA + 0.02% NaN_3_ for 2 h at room temperature. 75 μL of blocking buffer and 50 μL of hybridoma supernatant were combined, and 50 μL/well of diluted supernatants were added to the plates and incubated for 1 h at room temperature. Bound IgG was detected using HRP-conjugated goat anti-mouse IgG (at 1:2,000). Following a 1 h incubation, washed plates were developed with 50 μL of 1-Step Ultra TMB-ELISA, quenched with 2 N H_2_SO_4_, and the absorbance was read at 450nm.

### Competition binding analysis

The assay was performed as described previously (Zost *et al*., 2020). Briefly, for screening study wells of 384-well microtiter plates were coated with 1 μg/mL of purified SARS-CoV-2 S6P_ecto_ protein at 4 °C overnight. Plates were blocked with 2% bovine serum albumin (BSA) in DPBS-T for 1 h. Mouse hybridoma culture supernatants were diluted five-fold in blocking buffer, added to the wells (20 μl per well) in duplicates for each tested reference mAb and incubated for 1 h at room temperature. Biotinylated reference human mAbs with known epitope specificity (COV2-2130, COV2-2196 (Zost *et al*., 2020), and CR3022 (ter Meulen et al., 2006)) were added to each of well with the respective hybridoma culture supernatant at 1.25 μg/mL in a volume of 5 μl per well (final concentration of biotinylated mAb, 0.25 μg/mL) without washing of the plates, and then incubated for 1 h at room temperature. Plates then were washed, and bound antibodies were detected using HRP-conjugated avidin (Sigma, A3151, 0.3 μg/mL final concentration) and a TMB substrate. The signal obtained for binding of the biotin-labelled reference antibody in the presence of the hybridoma culture supernatant was expressed as a percentage of the binding of the reference antibody alone after subtracting the background signal. Tested mAbs were considered competing if their presence reduced the reference antibody binding to less than 41% of its maximal binding and non-competing if the signal was greater than 71%. A level of 40–70% was considered intermediate competition.

### Human ACE2 binding inhibition analysis

The assay was performed as described previously (Zost *et al*., 2020). Briefly, for screening study wells of 384-well microtiter plates were coated with 1 μg/mL purified recombinant SARS-CoV-2 S6P_ecto_ protein at 4 °C overnight. Plates were blocked with 2% non-fat dry milk and 2% normal goat serum in DPBS-T for 1 h. Mouse hybridoma culture supernatants were diluted five-fold in blocking buffer, added to the wells (20 μl per well) in quadruplicate, and incubated for 1 h at room temperature. Recombinant human ACE2 with a C-terminal Flag tag peptide was added to wells at 2 μg/mL in a 5 μl per well volume (final 0.4 μg/mL concentration of human ACE2) without washing of the plates, and then incubated for 40 min at room temperature. Plates were washed and bound human ACE2 was detected using HRP-conjugated anti-Flag antibody (Sigma-Aldrich, A8592, 1:5,000 dilution) and TMB substrate. ACE2 binding without antibody served as a control for maximal binding. Antibody COV2-2196 (RBD) served as a control for ACE2 binding inhibition. The signal obtained for binding of the human ACE2 in the presence of each dilution of tested culture supernatant was expressed as a percentage of the human ACE2 binding without antibody after subtracting the background signal.

### Sequencing, cloning, and expression of chimeric IgG1

To generate chimeric human IgG1 from mouse hybridoma cell lines, cells were lysed in Trizol (Thermo) followed by RNA purification with Direct-Zol Micro kit (Zymo). 5’ RACE products were generated with Template Switching RT Enzyme Mix (New England Biolabs) using anchored poly(dT)23 and TSO (GCT AAT CAT TGC AAG CAG TGG TAT CAA CGC AGA GTA CAT rGrGrG) oligonucleotides according to the manufactures instructions. Heavy and light chain sequences were amplified with primers specific for the TSO handle-sequence and the respective constant region sequence with Q5 Polymerase (New England Biolabs). Following Sanger sequencing, full-length variable regions were synthesized as gene blocks (Integrated DNA Technologies) and cloned into hIgG1 and hKappa expression vectors by Gibson assembly. Recombinant antibodies were expressed in Expi293 cells following co-transfection of heavy and light chain plasmids (1:1 ratio) using Expifectamine 293 (Thermo Fisher Scientific). Supernatants were harvested after 5-6 days, purified by affinity chromatography (Protein A Sepharose, GE), and desalted with a PD-10 (Cytiva) column.

### Binding analysis via biolayer interferometry

Biolayer interferometry (BLI) was used to quantify the binding capacity of SARS2-38 Fab fragments to trimerized SARS-CoV-2 spike. 10 µg/mL of biotinylated spike was immobilized onto streptavidin biosensors (ForteBio) for 3 min. After a 30 sec wash, the pins were submerged in running buffer (10 mM HEPES, 150 mM NaCl, 3 mM EDTA, 0.05% P20 surfactant, and 1% BSA) containing SARS2-38 Fab ranging from 1 to 1,000 nM, followed by a dissociation step in running buffer alone. The BLI signal was recorded and analyzed using BIAevaluation Software (Biacore).

### Cryo-EM sample preparation

Data were collected on lacey carbon grids with or without ultra-thin carbon film. For standard lacey carbon grids (Ted Pella #01895-F), SARS-CoV-2 spike was prepared at 1 mg/mL in TBS (30mM Tris pH 8, 150mM NaCl). For lacey carbon grids with ultra-thin carbon film (Ted Pella #01824G), SARS-CoV-2 spike was prepared at 0.2 mg/mL in TBS. Each sample was incubated for 15 min with 1 molar equivalent of SARS2-38 Fab fragments, applied to glow-discharged grids, then flash-frozen in liquid ethane using a Vitrobot Mark IV (ThermoFisher Scientific).

### Cryo-EM data collection

Grids were loaded into a Cs-corrected FEI Titan Krios 300kV microscope equipped with a Falcon 4 direct electron detector. Images were collected at a nominal magnification of 59000x, resulting in a pixel size of 1.16Å. Each movie consisted of 50 frames at 260ms each with a dose of 1e-/Å2/frame, yielding a total dose of 50e-/Å2/movie.

### Cryo-EM data processing

Movies were motion corrected using MotionCor2 v1.3.1 (Zheng et al., 2017), and contrast transfer function parameters were estimated using GCTF v1.18 (Zhang, 2016). Particles were picked using a general model in CrYOLO v1.7.6 (Wagner et al., 2019). 2D classification was performed in Relion 3.1 (Scheres, 2012; Zivanov et al., 2018), and particles in good classes from grids with or without ultra-thin carbon were combined for further processing. These particles were subjected to 3D classification, and those from the best class (all RBDs in the down position, with one bound by Fab) were selected for iterative Bayesian polishing and per-particle CTF refinement in Relion 3.1 (Zivanov et al., 2019). These particles were then used in non-uniform refinement in cryoSPARC v3.1.0 to generate a full-spike map (Punjani et al., 2017). To improve map quality at the Fab/spike interface, a mask was generated encompassing only the Fv and RBD, and particles were subjected to local non-uniform refinement in cryoSPARC v3.1.0. Final maps were sharpened via deep learning employed through DeepEMhancer (Sanchez-Garcia et al., 2020).

### Model building

The locally refined map was used to construct a model of the RBD bound by SARS2-38 Fv. An initial model for the RBD was adapted from a crystal structure of RBD bound to ACE2 (PDB 6M0J). For initial modeling of SARS2-38 Fv, pBLAST was used to identify pre-existing Fab structures with high sequence similarity (PDB 1KIQ for VH, and PDB 5XJM for VL). These starting components were combined and docked into the map, then refined in Coot v0.9.5 (Emsley et al., 2010), Isolde v1.1.0 (Croll, 2018), and Phenix v1.19 (Adams et al., 2010). Epitope and paratope contacts were identified using qtPISA (Krissinel and Henrick, 2007), and structures were visualized using UCSF ChimeraX (Goddard et al., 2018).

The full-spike map was used to construct a model of the spike bound by one Fv with all RBDs in the down position. An initial model was generated by combining the locally refined Fv/RBD structure with a previously solved cryo-EM structure of trimeric SARS-CoV-2 spike in the proper RBD configuration (PDB 6VXX). This model was docked into the full-spike map then refined using Coot v0.9.5, Isolde v1.1.0, and Phenix v1.19.

### RBD conservation analysis

RBD sequence data (residues 333-520) were retrieved on March 28, 2021 from the COVID-19 CoV Genetics Browser (covidcg.org), enabled by data from GISAID (Chen et al., 2021a; Shu and McCauley, 2017). In total, 786,273 sequences were included in the analysis. Probability of conservation relative to the reference sequence (2019n-CoV/WA1/2020) was computed for each residue, and results were log-transformed and normalized to generate a per-residue conservation score (1 = complete conservation, 0 = zero conservation). Results were visualized using a color-coded surface rendering of the RBD in UCSF ChimeraX.

### Neutralization assays

FRNTs were performed as described (Case *et al*., 2020). Briefly, serial dilutions of antibody were incubated with 2 x 10^2^ FFU of SARS-CoV-2 for 1 h at 37°C. Immune complexes were added to cell monolayers (Vero E6 cells or other cell lines where indicated) and incubated for 1 h at 37°C prior to the addition of 1% (w/v) methylcellulose in MEM. Following incubation for 30 h at 37°C, cells were fixed with 4% paraformaldehyde (PFA), permeabilized and stained for infection foci with SARS2-16 (hybridoma supernatant diluted 1:6,000 to a final concentration of ∼20 ng/mL) when using SARS-CoV-2 isolate WA1/2020, or with a mixture of mAbs that bind various epitopes on the RBD and NTD of spike (SARS2-02, SARS2-11, SARS2-31, SARS2-38, SARS2-57, and SARS2-71; diluted to 1 µg/mL total mAb concentration) for the VOCs. Antibody-dose response curves were analyzed using non-linear regression analysis (with a variable slope) (GraphPad Software). The antibody half-maximal inhibitory concentration (EC50) required to reduce infection was determined.

### Pre- and post-attachment neutralization assays

For pre-attachment assays, serial dilutions of mAbs were prepared at 4°C in Dulbecco’s modified Eagle medium (DMEM) with 2% FBS and preincubated with 10^2^ FFU of SARS-CoV-2 for 1 h at 4°C. MAb-virus complexes were added to a monolayer of Vero cells for 1 h at 4°C. Virus was allowed to internalize during a 37°C incubation for 30 min. Cells were overlaid with 1% (wt/vol) methylcellulose in MEM. For post-attachment assays, 2 x 10^2^ FFU of SARS-CoV-2 was adsorbed onto a monolayer of Vero cells for 1 h at 4°C. After removal of unbound virus, cells were washed twice with cold DMEM, followed by the addition of serial dilutions of MAbs in cold DMEM. Virus-adsorbed cells were incubated with mAd dilutions for 1 h at 4°C. Virus then was allowed to internalize for 30 min at 37°C, and subsequently cells were overlaid with methylcellulose as described above. Thirty hours later, plates were fixed with 4% PFA and analyzed for antigen-specific foci as described above for FRNTs.

### Attachment inhibition assay

SARS-COV-2 was incubated with mAbs at 10 µg/mL for 1 h at 4°C. The mixture then was added to pre-chilled Vero E6, Vero-TMPRSS2, Vero-TMPRSS2-ACE2, or Calu-3 cells at an MOI of 0.005 and incubated at 4°C for 1 h. Cells were washed six times with chilled PBS before addition of lysis buffer and extraction of RNA using MagMax viral RNA isolation kit (Thermo Fisher Scientific) and a Kingfisher Flex 96-well extraction machine (Thermo Fisher Scientific). SARS-CoV-2 RNA was quantified by qRT-PCR using the N-specific primer/probe set described below. *GAPDH* was measured using a predesigned primer/probe set (IDT PrimeTime Assay Hs.PT.39a.22214836). Viral RNA levels were normalized to GAPDH, and the fold change was compared with isotype control mAb. For each cell type, a control with a 4-fold lower MOI (0.00125) was included to demonstrate detection of decreased viral RNA levels.

### Virus internalization assay

SARS-COV-2 was incubated with mAbs at 10 µg/mL for 1 h at 4°C. The mixture was then added to pre-chilled Vero E6 cells at an MOI of 0.005 and incubated at 4°C for 1 h. Cells were washed twice with chilled PBS to remove unbound virus, and subsequently incubated in DMEM at 37°C for 30 min to allow virus internalization. Cells then were treated with proteinase K and RNaseA at 37°C for 10 min to removed uninternalized virus. Viral and cellular RNA were extracted and analyzed as described above for the attachment inhibition assay. A no internalization control was included, where proteinase K and RNase A treatments were performed directly after washing, without an internalization step.

### Measurement of viral burden and cytokine and chemokine levels

On 7 dpi, mice were euthanized and organs were collected. Nasal washes were collected in 0.5 mL of PBS. Organs were weighed and homogenized using a MagNA Lyser (Roche). Viral RNA from homogenized organs or nasal wash was isolated using the MagMAX Viral RNA Isolation Kit (ThermoFisher) and measured by TaqMan one-step quantitative reverse-transcription PCR (RT-qPCR) on an ABI 7500 Fast Instrument. Viral burden is expressed on a log_10_ scale as viral RNA per mg for each organ or total nasal wash after comparison with a standard curve produced using serial 10-fold dilutions of viral RNA standard. Primers were 5’-ATGCTGCAATCGTGCTACAA-3’, 5’-GACTGCCGCCTCTGCTC-3’, and probe 5’-/56-FAM/ TCAAGGAAC/Zen/ AACATTGCCAA/3IABkFQ-3’ (Case *et al*., 2020). For the measurement of cytokine and chemokine levels in the lung, lung homogenates were treated with 1% Triton X-100 for 1 h at room temperature to inactivate virus. Cytokine and chemokine levels in the lung homogenate were then analyzed by multiplex array (Eve Technologies Corporation).

## QUANTIFICATION AND STATISTICAL ANALYSIS

Statistical significance was assigned when p values were < 0.05 using Prism version 8 (GraphPad). Tests, number of animals (n), median values, and statistical comparison groups are indicated in the Figure legends.

