## Supplemental Figures and Table for "A potently neutralizing anti-SARS-CoV-2 antibody inhibits variants of concern by binding a highly conserved epitope"

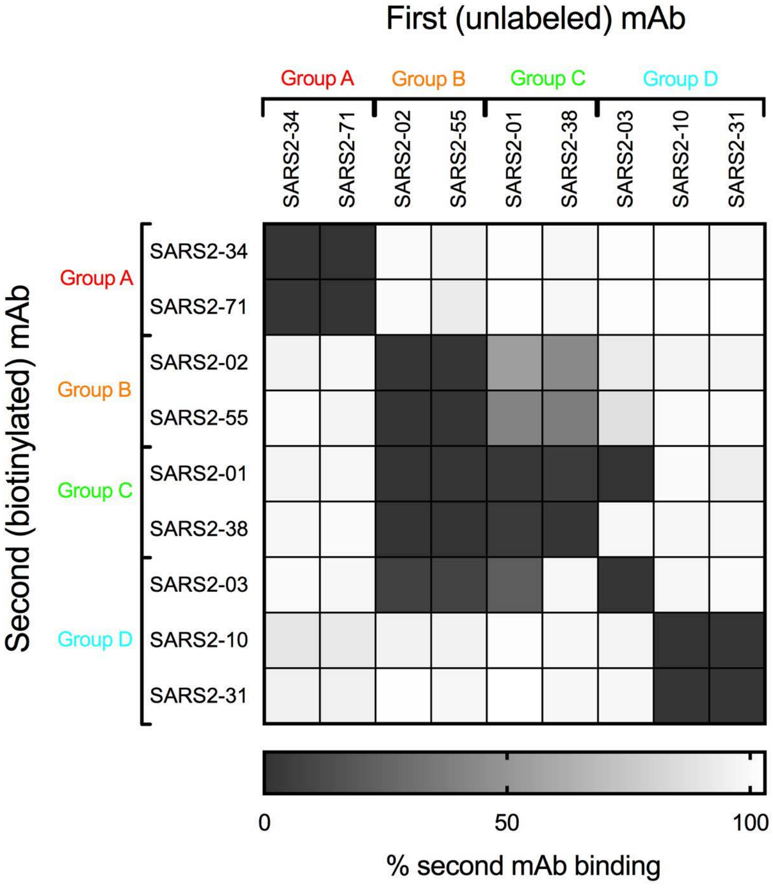
First (unlabeled) mAb
Group A
Group B
Group C
Group D
SARS2-34
SARS2-71
SARS2-02
SARS2-55
SARS2-01
SARS2-38
SARS2-03
SARS2-10
SARS2-31
Second (biotinylated) mAb
0
50
100
% second mAb binding

### Figure S1

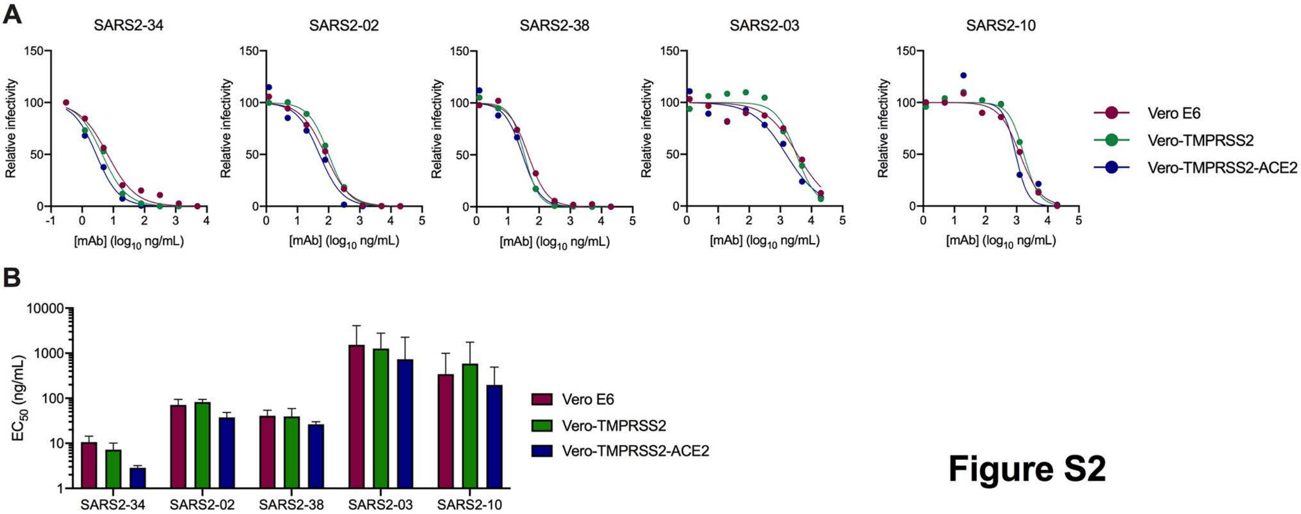
A
SARS2-34
SARS2-02
SARS2-38
SARS2-03
SARS2-10
Relative infectivity
[mAb] (log10 ng/mL)
Vero E6
Vero-TMPRSS2
Vero-TMPRSS2-ACE2
B
EC50 (ng/mL)
SARS2-34
SARS2-02
SARS2-38
SARS2-03
SARS2-10
Vero E6
Vero-TMPRSS2
Vero-TMPRSS2-ACE2

**Figure S2**

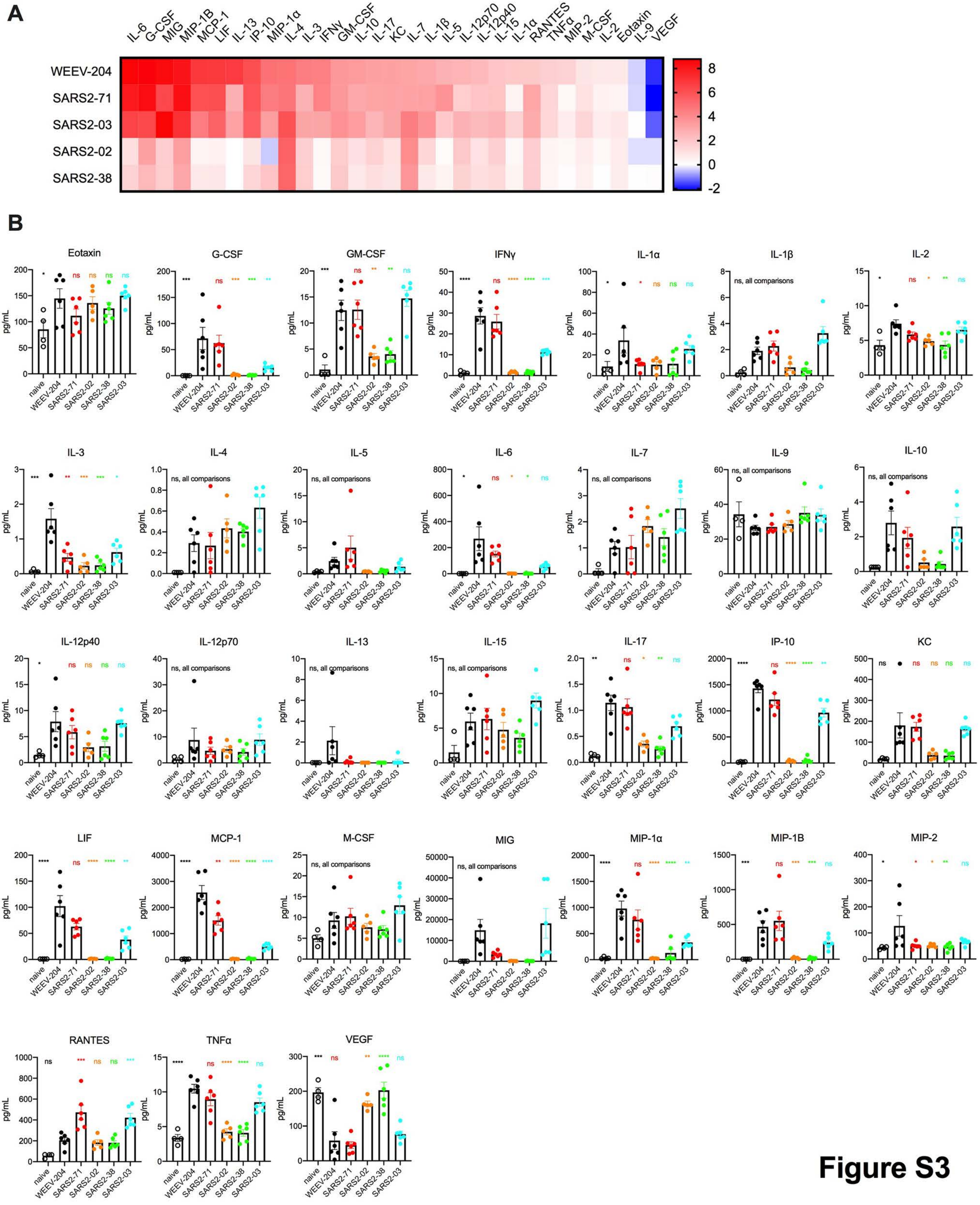
A
IL-6 G-CSF MIG MIP-1B MCP-1 LIF IL-13 IP-10 MIP-1α IL-4 IL-3 IFNγ GM-CSF IL-10 IL-17 KC IL-7 IL-1β IL-5 IL-12p70 IL-12p40 IL-15 IL-1α RANTES TNFα MIP-2 M-CSF IL-2 Eotaxin IL-9 VEGF
WEEV-204
SARS2-71
SARS2-03
SARS2-02
SARS2-38
8
6
4
2
0
-2
B
Eotaxin
G-CSF
GM-CSF
IFNγ
IL-1α
IL-1β
IL-2
IL-3
IL-4
IL-5
IL-6
IL-7
IL-9
IL-10
IL-12p40
IL-12p70
IL-13
IL-15
IL-17
IP-10
KC
LIF
MCP-1
M-CSF
MIG
MIP-1α
MIP-1B
MIP-2
RANTES
TNFα
VEGF
pg/mL
naive
WEEV-204
SARS2-71
SARS2-02
SARS2-38
SARS2-03
ns, all comparisons

Figure S3

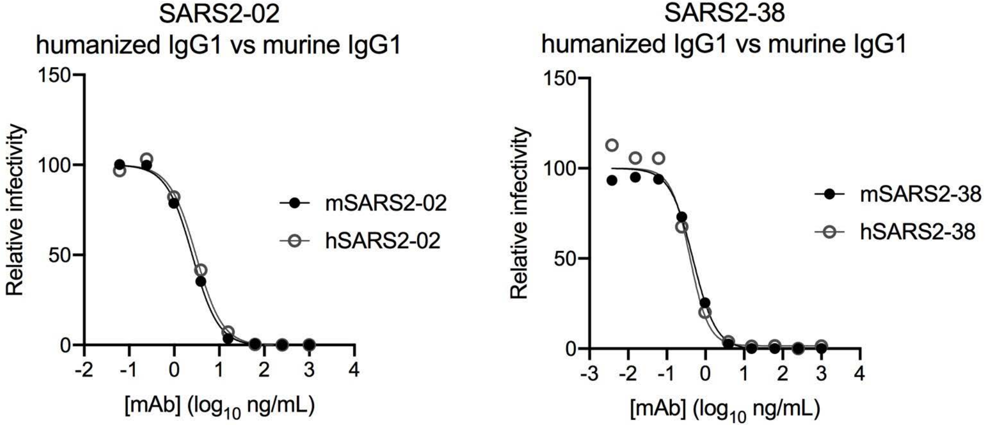
SARS2-02
humanized IgG1 vs murine IgG1
Relative infectivity
150
100
50
0
-2
-1
0
1
2
3
4
[mAb] (log10 ng/mL)
mSARS2-02
hSARS2-02
SARS2-38
humanized IgG1 vs murine IgG1
Relative infectivity
150
100
50
0
-3
-2
-1
0
1
2
3
4
[mAb] (log10 ng/mL)
mSARS2-38
hSARS2-38

### Figure S4

**Figure S5**

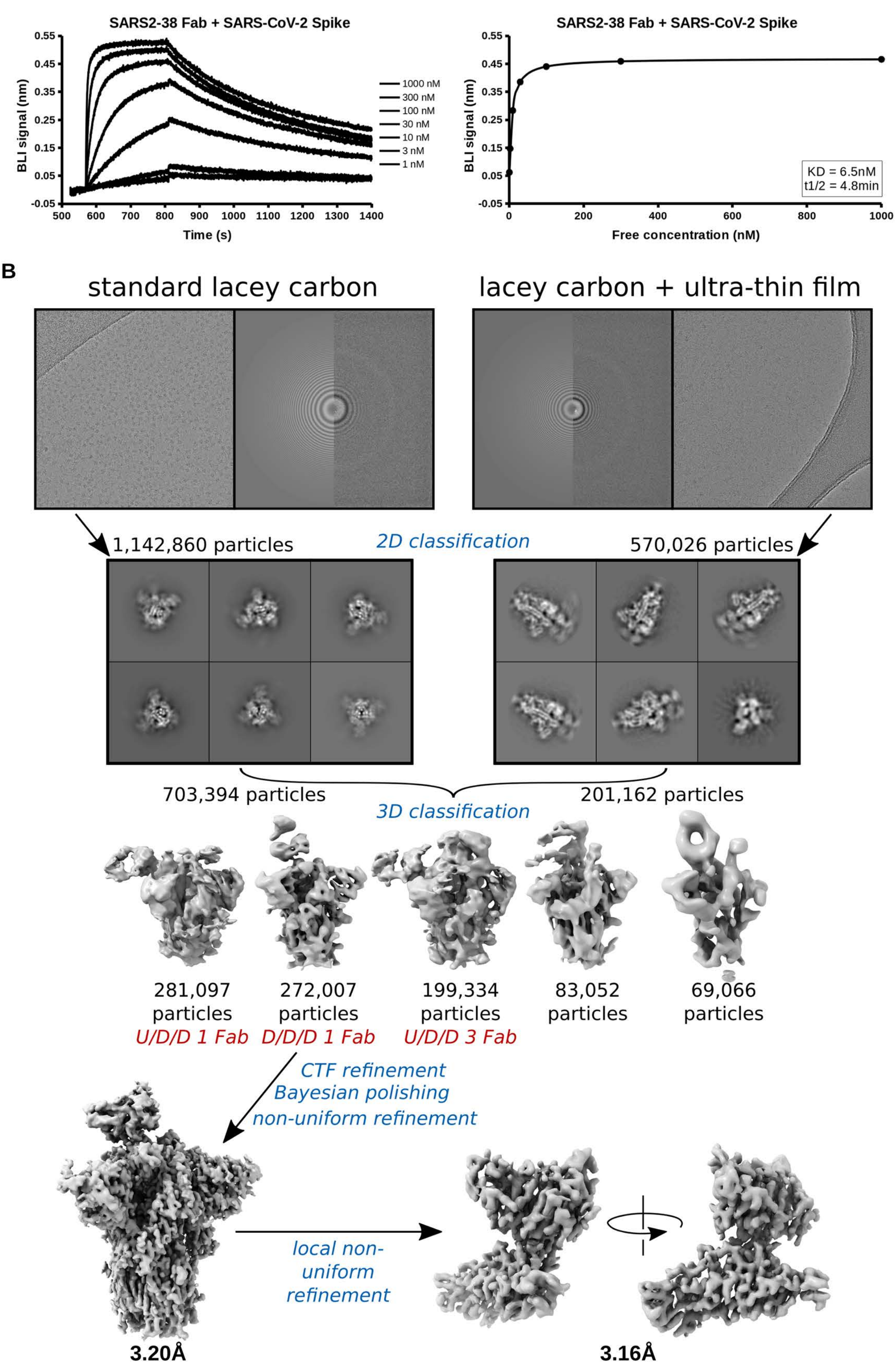
A
SARS2-38 Fab + SARS-CoV-2 Spike
BLI signal (nm)
Time (s)
1000 nM
300 nM
100 nM
30 nM
10 nM
3 nM
1 nM
SARS2-38 Fab + SARS-CoV-2 Spike
BLI signal (nm)
Free concentration (nM)
KD = 6.5nM
t1/2 = 4.8min
B
standard lacey carbon
lacey carbon + ultra-thin film
1,142,860 particles
2D classification
570,026 particles
703,394 particles
201,162 particles
3D classification
281,097 particles
U/D/D 1 Fab
272,007 particles
D/D/D 1 Fab
199,334 particles
U/D/D 3 Fab
83,052 particles
69,066 particles
CTF refinement
Bayesian polishing
non-uniform refinement
local non-uniform refinement
3.20Å
3.16Å

#### Figure S6

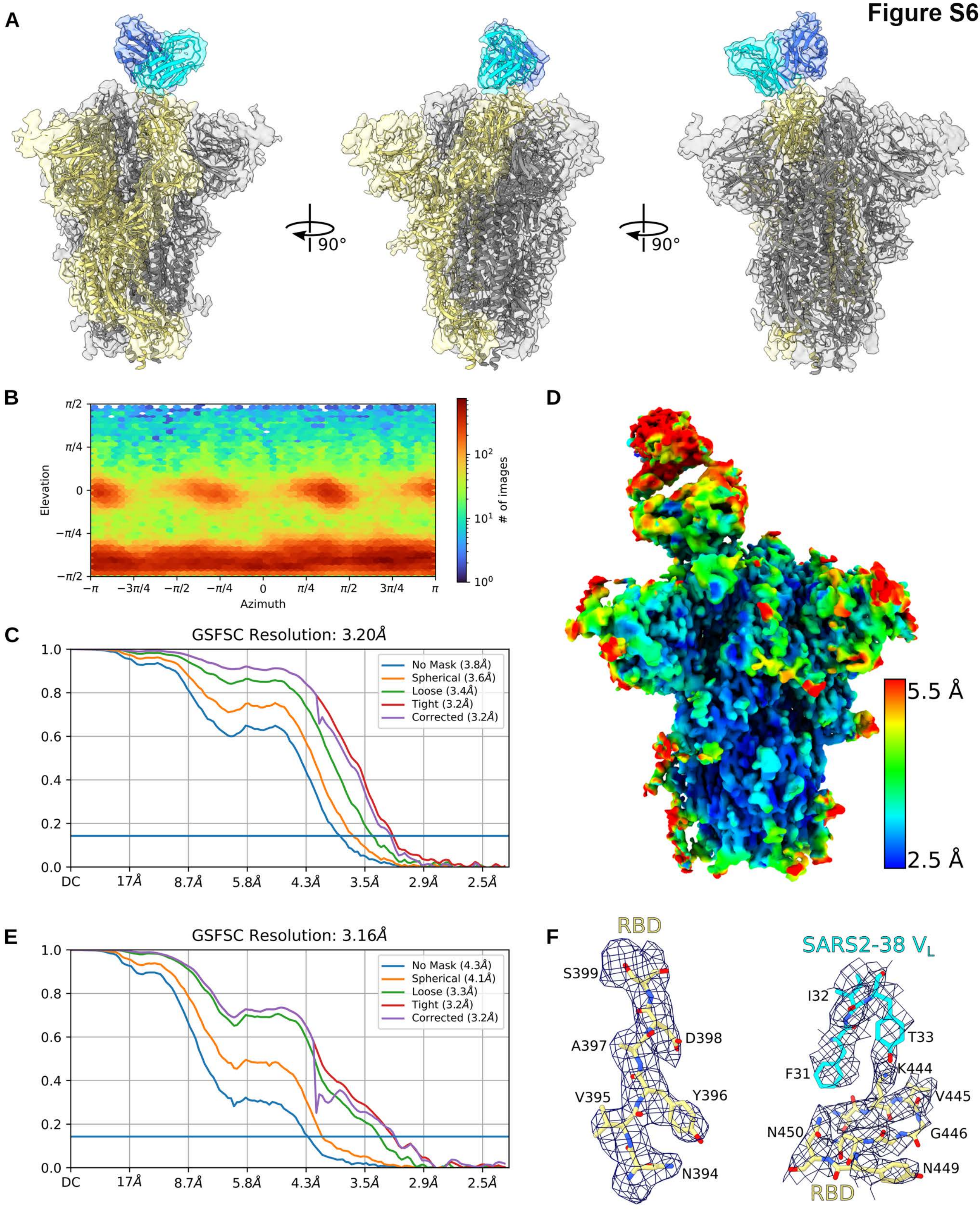
A
90°
90°
B
π/2
π/4
0
−π/4
−π/2
Elevation
−π
−3π/4
−π/2
−π/4
0
π/4
π/2
3π/4
π
Azimuth
$10^2$
$10^1$
$10^0$
### of images
C
GSFSC Resolution: 3.20Å
No Mask (3.8Å)
Spherical (3.6Å)
Loose (3.4Å)
Tight (3.2Å)
Corrected (3.2Å)
DC
17Å
8.7Å
5.8Å
4.3Å
3.5Å
2.9Å
2.5Å
D
5.5 Å
2.5 Å
E
GSFSC Resolution: 3.16Å
No Mask (4.3Å)
Spherical (4.1Å)
Loose (3.3Å)
Tight (3.2Å)
Corrected (3.2Å)
F
RBD
S399
A397
D398
V395
Y396
N394
SARS2-38 $V_L$
I32
T33
F31
K444
V445
N450
G446
N449
RBD

### Table S1

| Cryo-EM data collection, processing, and model refinement statistics | | |
|---|---|---|
| | SARS-CoV-2 spike + Fab SARS2-38 (global) PDB 7MKL EMD-23898 | SARS-CoV-2 spike + Fab SARS2-38 (local) PDB 7MKM EMD-23899 |
| **Data collection** | | |
| Magnification | 59,000x | 59,000x |
| Exposure ($e^-/Å^2$) | 50 | 50 |
| Defocus range (μm) | 0.8-2.3 | 0.8-2.3 |
| Pixel size (Å/pixel) | 1.16 | 1.16 |
| **Data processing** | | |
| Initial particles (no.) | 1,712,886 | 1,712,886 |
| Final particles (no.) | 272,007 | 272,007 |
| Nominal resolution (Å) | 3.20 | 3.16 |
| FSC threshold | 0.143 | 0.143 |
| **Model refinement** | | |
| Adapted PDB models | 6M0J, 1KIQ, 5XJM, 6VXX | 6M0J, 1KIQ, 5XJM |
| Model resolution (Å) | 4.1 | 4.2 |
| FSC threshold | 0.5 | 0.5 |
| Model composition | | |
| Non-hydrogen atoms | 25,597 | 3,192 |
| Residues | 3,164 | 406 |
| Ligands | 63 | 1 |
| B-factors ($Å^2$) | | |
| Residues | 102.17 | 76.19 |
| Ligands (glycans) | 122.20 | 81.01 |
| Bonds (RMSD) | | |
| Length (Å) | 0.004 | 0.003 |
| Angles (°) | 0.816 | 0.797 |
| Validation | | |
| Molprobity score | 1.69 | 1.51 |
| Clash score | 10.10 | 8.33 |
| Rotamer outliers (%) | 0.00 | 0.00 |
| Ramachandran | | |
| Favored (%) | 97.09 | 97.75 |
| Allowed (%) | 2.91 | 2.25 |
| Outliers (%) | 0.0 | 0.0 |
